# Regional brain age deviations reveal divergent developmental pathways in youth

**DOI:** 10.1101/2025.11.27.690886

**Authors:** Anthony Gagnon, Marie A. Brunet, Maxime Descoteaux, Larissa Takser

## Abstract

**Background:** Normative modeling of brain development has gained traction for quantifying individual deviations in maturation. The brain age gap (BAG), the difference between predicted age from MRI features and chronological age, offers a potential individualized normative metric of neurodevelopment. However, consistent patterns across psychiatric disorders remain elusive, and no studies have examined whether BAG can predict developmental trajectories within an inclusive continuous model of youth’s cognition and behavior.

**Methods:** Using longitudinal data from the Adolescent Brain Cognitive Development Study (ages 9-15, n=9,074), we built 8 region-specific brain age models using volumes, thicknesses, and surface areas of parcels from the Brainnetome adolescent atlas. We derived psychiatric diagnoses from a parental questionnaire. Multivariate linear regression was used to assess case-control differences and cross-sectional continuous cognitive/behavioral profiles. We modeled cognitive/behavioral trajectories using a multivariate joint latent-class mixed model and assessed the relationship with BAG values using multinomial logistic regression.

**Results:** Children with ADHD showed delayed maturation across multiple regions (Cohen’s *d*: - 0.12 to -0.08), while subcortical BAG emerged as a transdiagnostic indicator of delayed development (*d*: -0.07, *p_fdr_* = 0.024). Accelerated maturation characterized the high cognition and low symptom profile, while the inverse was found for the low cognition profile. Three developmental trajectories were identified: stable, towards externalizing behaviors, or internalizing behaviors. Widespread accelerated maturation predicted evolution towards internalizing behaviors but was protective against the externalizing trajectory.

**Conclusions:** Integrating BAG with continuous cognitive and behavioral profiles yielded a plausible framework for early identification of atypical trajectories, potentially contributing to personalized medicine in psychiatry.

## Introduction

Normative modeling has gained traction in neuroimaging in recent years, aiming to provide informative brain charts for use in clinical care(1). Those charts provide valuable information on the neurotypical trajectories of brain maturation, especially during childhood and adolescence, when both myelination and pruning occur concurrently throughout the brain(2). One primary aim of these charts is to provide a clinical tool for assessing and identifying individual divergent brain developmental trajectories. Such clinical tools are of utmost importance in childhood/adolescent psychiatry, where symptom presentations are complex and require the inclusion of an ever-increasing number of data sources(3). Unsurprisingly, those developmental periods coincide with the onset of many psychiatric disorders(4,5), representing a substantial societal burden worldwide(6).

In the past decade, the brain age gap (BAG) framework has been proposed as an individualized normative metric of brain developmental stages(7–9). Conceptually, this metric relies on training a model to predict chronological age from brain features derived from MRI acquisitions. Computing the difference between the model’s predicted age and the chronological age results in a quantitative assessment of brain trajectories. In pediatric populations, individuals with negative BAG values are thought to exhibit delayed brain development (e.g., delayed maturation), whereas those with positive BAG values are considered to have an accelerated developmental trajectory (e.g., accelerated maturation). Applications of this framework to understand either delayed or accelerated development in pediatric psychiatric diagnoses and psychopathology show mixed results. Indeed, studies showed higher BAG values in clinical populations with high risks for psychosis(10). Oppositely, lower BAG values in autism spectrum disorder, attention deficit-hyperactivity disorder (ADHD), or intellectual disability were also reported(11–13). Researchers noted an association between delayed development, as indicated by lower BAG values, and higher psychopathology symptoms in two large pediatric cohorts(14,15). Those results place the BAG indicator as an interesting clinical biomarker of psychiatric disorders and psychopathology; however, its understanding remains limited and warrants future studies. Additionally, most studies focused on cross-sectional associations and did not investigate whether BAG could become a potential biomarker of developmental trajectories. Given the current need in precision psychiatry(16,17), a promising biomarker must provide information about future developmental trajectories to facilitate early identification and guide efficient interventions.

Aside from psychopathology, the BAG framework has also been applied to understand the spread of cognitive abilities found in the general population. While a positive relationship between BAG and various cognitive functions (e.g., working memory, arithmetic skills, or a more general cognitive score) has been reported(18–21), others have found no significant relationship(22,23). However, most of those studies considered either psychopathology or cognition as two separate entities, even though there exists a relationship between behavioral manifestations and cognitive abilities(24–26). We recently proposed a new fuzzy framework encompassing four profiles that represent the continuum of behavioral manifestations and cognitive abilities in general pediatric populations(27). One crucial advantage of this framework, compared to other clustering techniques, is its ability to quantify the extent to which a single participant identifies with each profile through membership values. In practical terms, this means participants identify with each profile along a continuous axis, aligning with recent domain-based frameworks(28,29), which is more suitable for real-world applications than discrete classification. Additionally, we demonstrated the longitudinal reproducibility of the profiles in subsequent follow-ups of the Adolescent Brain Cognitive Development cohort(27,30). Putting this altogether, this profiling framework represents a unique opportunity to evaluate the relationship between BAG and cognition/behavior when assessed jointly. Additionally, the ability to track movement through profiles during development raises the question of whether the BAG at baseline can predict future cognitive and behavioral trajectories. Investigating this relationship would provide valuable insights into BAG’s potential as a biomarker, as predicting future trajectories is of utmost importance in precision psychiatry.

By leveraging longitudinal data from one of the largest pediatric neuroimaging cohorts, ABCD(30,31), we built whole-brain and regional brain age models using a pediatric-tailored brain atlas(32). Using the derived BAG values for each region, we evaluated whether BAG values differed between participants with a psychiatric diagnosis and typically developing (TD) participants. Then, leveraging our previously proposed fuzzy cognitive and behavioral profiles(27), we assessed whether BAG values would be associated with the baseline profiles’ membership values when cognition and behavior were jointly considered. Finally, we modeled trajectories within our fuzzy profiles framework and assessed whether baseline BAG values could predict specific developmental trajectories. For all aims, we hypothesize that delayed development, marked by lower BAG values, would be associated with a negative outcome.

## Methods and Materials

### Participants

Primary analyses were conducted on participants with available baseline structural MRI, cognitive, and behavioral assessments drawn from the Adolescent Brain Cognitive Development (ABCD) Study Release 5.1(30). The ABCD Study is a large-scale, multi-site, longitudinal, and prospective pediatric cohort comprising 11,878 children recruited through the U.S. school system(30). Initial enrollment occurred between 2016 and 2018 during the 9- to 11-year follow-up period. Additionally, participants who completed the subsequent 11-13y and 13-15y follow-ups with available cognitive and behavioral data, as described elsewhere(27), were included in secondary analyses assessing the developmental trajectories. For more details regarding data releases, participating sites, ethics, study protocols, and investigators, please see https://docs.abcdstudy.org/latest/.

### Image processing, quality control, and harmonization

T1-weighted images were acquired for all participants using a harmonized protocol across the 21 participating sites(31). As in a previous study(33), participants whose MRI acquisitions were performed on a Philips scanner were excluded to facilitate post-hoc harmonization across sites. Acquisition parameters between the remaining GE and Siemens scanners were nearly identical (FOV: 256 x 256, 1 mm isotropic, TR: 2500 ms, and TE: 2.88 ms (Siemens) or 2 ms (GE))(31). Raw images that passed initial quality control by the ABCD Study team were preprocessed, segmented, and converted into surfaces using the FastSurfer(34) deep-learning tool via the nf-pediatric pipeline (https://github.com/scilus/nf-pediatric)(35–38). To ensure age-appropriate cortical and subcortical segmentations, we mapped the newly proposed preadolescent Brainnetome atlas (227 cortical and subcortical regions) using surface-based registration in the subject space(32). Using surface-based registration rather than volumetric registration yields better alignment with individual cortical folding patterns(39). Following surface-based registration, metrics, including surface area, thickness, and volume, were computed for each cortical parcel using FreeSurfer(40). For subcortical parcels, only volume was calculated and used in the brain age models, while all three metrics were used for cortical parcels.

Quality control for each subject was performed based on two criteria: 1) all cortical/subcortical segmentations were successfully extracted, and 2) showed metric values within five times the interquartile range (± 5 x *IQR*) for each parcel. This statistical approach ensures effective, robust quality control across a large number of subjects. We selected 5 times the interquartile range rather than the classical 3 times to ensure we removed only true outliers and not scanner or site artifacts, since this quality control step was performed before harmonization across sites. A flowchart of participants’ exclusion, along with their associated reasons, is shown in Supplementary Figure 1. Detailed charts with the distribution for each parcel and metric can be seen in Supplementary Figures 2, 3, 4, 5, 6, and 7.

Harmonization across sites was performed using a pairwise version of ComBat(41,42). The pairwise approach requires selecting a reference site, to which all the other sites will be harmonized. We selected site #16 as our reference site due to its large number of participants and sex/age distribution matching the overall distribution found in ABCD (Supplementary Figure 8). Harmonization was then performed for each brain region and across measures (volume, thickness, and surface area) while retaining the variance explained by age, sex, and handedness. Example results for a single brain region are presented in Supplementary Figures 9, 10, and 11.

### Brain age models

We built 24 brain age models using the eXtreme Gradient Boosting framework(43). First, we trained a model to predict brain age on the entire brain, comprising 227 parcellations. Second, we trained individual models for each of the following coarse brain regions: subcortical/cerebellar, frontal, temporal, parietal, insular, limbic, and occipital areas, resulting in a total of eight models, as previously done by Kaufmann *et al.* (2019)(44) (Figure 1). Included features for each model are detailed in Supplementary Table 1. To assess sex differences and perform sensitivity analyses, we then repeated the process for both males and females, yielding 24 brain age models in total. Throughout this manuscript, the use of “male” and “female” refers to the biological sex of the participants, and in no case assumes their gender.

**Figure 1.**
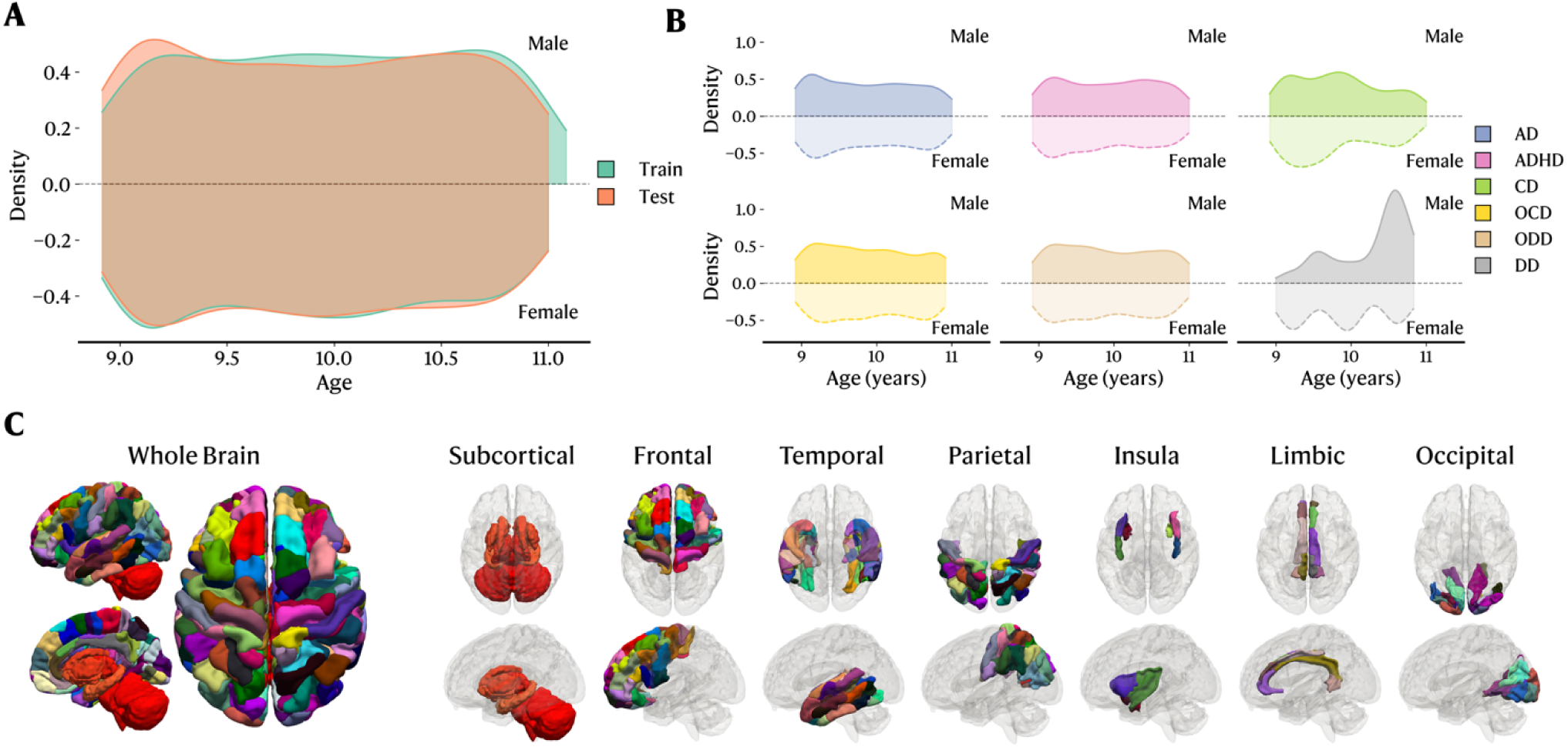
Sample distribution across training, testing, and diagnosis sets, combined with brain regions included in all eight brain age models. **A.** Age distribution, stratified by sex, between the train and test sets. Both training and testing samples include only TD hildren. **B.** Age distribution stratified by sex for the hold-out (not included in training/testing) clinical sample. **C.** Brain features ncluded in all eight brain age models. AD: Anxiety disorder. ADHD: Attention-deficit/hyperactivity disorder. CD: Conduct disorder. OCD: Obsessive-compulsive disorder. ODD: Oppositional defiant disorder. DD: Depressive disorder. All psychopathology diagnoses re derived from the KSADS and do not reflect a clinical diagnosis.

Consistent with previous literature(44), we left out all participants with a clinical diagnosis of psychiatric disorders and performed model training solely on TD participants (Figure 1). The final prediction was performed using the entire sample. For each model (except the individual male/female models, which used a 60/40 training/testing split), the remaining 50% of subjects were included in the training set, while the other half was kept as the testing set. As performed in previous studies(45,46), we ensured that participants from the same family were not split between the training and testing sets. Hyperparameters were then tuned using Bayesian optimization with 5-fold cross-validation over a wide range of parameter values, and the resulting optimal parameters were used to fit the final model on the training set. The fitted models were then evaluated on the test set using the mean absolute error (MAE) and Pearson correlation coefficient (*r*). The predicted brain ages were then used to calculate the BAG using equation (1).

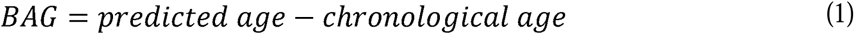

Brain age models are subject to age-related estimation biases that are commonly reported and discussed in the literature(47–52). Briefly, younger participants’ ages are systematically overestimated, whereas older participants’ ages are consistently underestimated. Therefore, we applied a correction method developed by Beheshti *et al.* (2019)^28^, which involves fitting a linear model of BAG against chronological age (Ω) in the training set to calculate an offset (ϕ) using equation (2). The choice of correction methods does not influence downstream results, nor does it alter the results compared to those of non-corrected BAG when also correcting for age(47). This offset was then subtracted from the predicted brain age to remove the age bias from the prediction (Supplementary Figure 12).

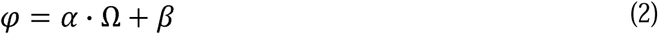

### Psychopathology, cognitive, and behavioral data

Psychiatric diagnoses were evaluated using the parent version of the Kiddie Schedule for Affective Disorders and Schizophrenia for School-Aged Children (KSADS) during the 9-11y follow-up. Based on prior work(53), we extracted six categorical diagnosis variables: anxiety disorder (AD), attention deficit-hyperactivity disorder (ADHD), conduct disorder (CD), depressive disorder (DD), obsessive-compulsive disorder (OCD), and oppositional defiant disorder (ODD). Additionally, we created a composite psychopathology index (PSYPATHO), which regroups participants with at least one of the above-mentioned psychiatric diagnoses. While we employ the term “psychiatric diagnoses” to refer to these categorical variables, it is important to note that they do not reflect clinical diagnoses, as they were not assessed by a clinician. Throughout this article, we will employ the term “suspected for” to refer to KSADS-derived diagnoses.

The complete method used to extract the cognitive and behavioral profiles has been extensively detailed elsewhere(27) and will only be briefly summarized here. During the 9- to 11-year follow-up, participants were administered the NIH Toolbox (NIHTB)(54), the Little Man’s Task (LMT)(55), the Rey Auditory Verbal Learning Test (RAVLT)(56), and the Wechsler Intelligence Test for Children-V Matrix Reasoning task(57). Participants were re-administered the NIHTB(54), LMT(55), and RAVLT(56) during the 11- to 13-year follow-up, and the NIHTB(54) and LMT(55) during the 13- to 15-year follow-up. For each follow-up, factor analysis was performed to extract latent scores for verbal ability (VA), executive function/processing speed (EFPS), and memory (MEM)(25). Those scores were combined with three behavioral symptom scores (internalizing, externalizing, and stress problems) from the Child Behavior Checklist (CBCL)(58). Both cognitive and behavioral scores were then harmonized across sites, residualized for covariates (age, sex, ethnicity, and handedness), and input into a fuzzy C-Means clustering algorithm, yielding a four-profile solution representing either high (HC/LB) or low (LC/LB) cognitive scores with low behavioral symptoms or high stress/internalizing behavior (MC/HSI) or high externalizing behavior (MC/HE) with moderate cognitive scores. The profiling process was performed independently for each follow-up, but returned a nearly identical profile structure(27). Membership values for all four profiles (e.g., the degree to which a participant aligns with each profile, ranging from 0 to 1) were extracted for each participant and used in subsequent statistical analyses.

### Statistical analyses

To investigate group effects between TD children and clinical samples, we applied multivariate linear regression, controlling for age, sex, handedness, ethnicity, parental education, and family income. For missing covariates, median imputation was used. We converted the t-statistics (*t*) into Cohen’s *d* effect sizes using equation (59), where *n*_1_ and *n*_2_ represent the number of participants in both groups. To assess the statistical significance of the effect sizes, we performed permutation testing by shuffling the diagnosis label across participants while keeping other covariates constant for 10,000 iterations. The resulting permutated p-value (*p_perm_*) was computed using equation (60), where *d^*(i)^*is the test statistic under the i^th^ permutation and *N* is the number of iterations performed. For all comparisons, clinical samples were compared only to those of TD children, and false discovery rate (FDR) correction was applied to all permuted p-values(61).

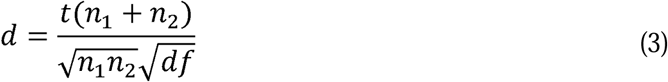

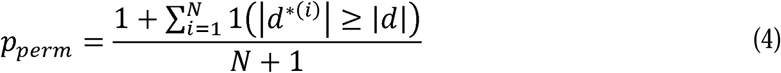

To assess the relationship between the BAG and baseline cognitive and behavioral profiles membership values, we applied a multivariate linear regression using the same covariates as previously mentioned. We converted the t-statistics (*t*) for the BAG into a partial correlation coefficient (*r_partial_*) using equation (59), reflecting the correlation attributable only to the BAG. Coefficient significance was also assessed using permutation testing with 10,000 iterations as described.

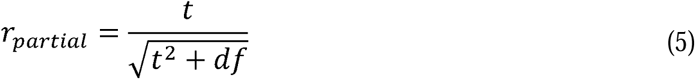

Longitudinal trajectories of profile membership values were modeled using a multivariate joint latent class mixed model from the *lcmm* R package(62). Briefly, for each cognitive/behavioral profile *k* ∈ {1, …, 4}, membership values (*Y*), subject *j*, repeated measurement *i*, and unobserved latent class *c_i_* ∈ {1, …, *G*}, we modeled the latent trajectories as described in equation (6) using class-independent fixed effects (*β_k0_*, *β_k1_*), class-specific slope (r_kg_), and residual error (∈_kij_). Then, the observed membership values are modeled using a 5-quantile splines monotone link function (*H*) corresponding to equation (7). This enables us to capture distinct developmental trajectories flexibly while also accounting for individual baseline starting points and rates of change. We fitted models for up to 10 classes using a grid search on starting values to avoid local optima. We selected the best model based on the entropy value and convergence status (Supplementary Figure 13).

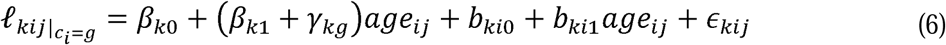

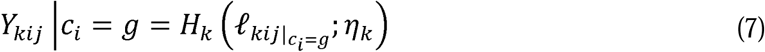

Following latent class fitting, we examined the relationship between BAG values and trajectories using secondary multinomial logistic regression models, using the stable class as reference, controlling for sex, ethnicity, handedness, parental education, and family income. Log-Odds Ratios were then transformed into Odds Ratios (OR) for easier interpretation. Where applicable, sensitivity analyses were conducted using male-only and female-only BAG models, with the same covariates and statistical methods as those used for the combined BAG model, except that the sex variable was excluded. All Python and R code to reproduce the analyses is available at https://github.com/gagnonanthony/Gagnon_BrainAge_2025.

## Results

Following quality control, 9,076 participants from the 9-11-year follow-up were included and used to build the brain age models. Cognitive and behavioral profile data from 7,368 and 2,845 participants were included from the 11-13-year and 13-15-year follow-up, respectively. Retaining only participants with all available time points, we included 1,897 participants for the longitudinal analysis of trajectories, while the remaining analyses were performed on the whole sample. Demographic information is available in Table 1 and Figure 1.

**Table 1.** Demographics Table for all follow-ups. GED: General Equivalent Diploma. AD: Anxiety Disorder. ADHD: Attention Deficit-Hyperactivity Disorder. CD: Conduct Disorder. DD: Depression Disorder. OCD: Obsessive-Compulsive Disorder. ODD: Oppositional Defiant Disorder. Unavailable data or empty groups are marked with a hyphen. All psychopathology diagnoses are derived from the KSADS and do not reflect a clinical diagnosis.

|  | 9-11y (Baseline) |  | 11-13y |  | 13-15y |  |
| --- | --- | --- | --- | --- | --- | --- |
|  | Male | Female | Male | Female | Male | Female |
| **N** | 4722 (52.04) | 4352 (47.96) | 3839 (52.1) | 3529 (47.89) | 1481 (52.04) | 1364 (47.93) |
| **Age (months)** | 119.22 (7.55) | 118.87 (7.5) | 143.58 (7.77) | 143.28 (7.76) | 168.93 (8.47) | 169.0 (8.3) |
| **Race/Ethnicity, count (%)** |  |  |  |  |  |  |
| White | 2443 (26.92) | 2152 (23.72) | 2176 (29.53) | 1948 (26.44) | 841 (29.55) | 743 (26.11) |
| Black or African American | 676 (7.45) | 650 (7.16) | 450 (6.11) | 455 (6.17) | 146 (5.13) | 162 (5.69) |
| Hispanic or Latino | 877 (9.66) | 851 (9.38) | 740 (10.04) | 679 (9.21) | 320 (11.24) | 269 (9.45) |
| Asian | 82 (0.9) | 89 (0.98) | 80 (1.09) | 77 (1.04) | 34 (1.19) | 31 (1.09) |
| Other | 487 (5.37) | 445 (4.9) | 393 (5.33) | 370 (5.02) | 140 (4.92) | 159 (5.59) |
| **Highest parental education, count (%)** |  |  |  |  |  |  |
| No Highschool | 176 (1.94) | 219 (2.41) | 161 (2.18) | 158 (2.14) | 63 (2.21) | 65 (2.28) |
| Highschool, GED, or equivalent | 426 (4.69) | 393 (4.33) | 319 (4.33) | 277 (3.76) | 108 (3.79) | 93 (3.27) |
| Some college | 1289 (14.21) | 1130 (12.45) | 951 (12.91) | 878 (11.91) | 373 (13.11) | 339 (11.91) |
| Bachelor's Degree | 1238 (13.64) | 1110 (12.23) | 1055 (14.32) | 949 (12.88) | 414 (14.55) | 367 (12.9) |
| Postgraduate Degree | 1583 (17.45) | 1495 (16.48) | 1353 (18.36) | 1267 (17.19) | 523 (18.38) | 500 (17.57) |
| **Familial Income (USD\$), count (%)** | | | | | | |
| < 50 000\$ | 877 (9.66) | 826 (9.1) | 660 (8.96) | 596 (8.09) | 250 (8.78) | 215 (7.55) |
| 50 000 - 100 000\$ | 1598 (17.61) | 1503 (16.56) | 1326 (17.99) | 1276 (17.32) | 515 (18.1) | 495 (17.39) |
| > 100 000\$ | 1848 (20.37) | 1674 (18.45) | 1554 (21.09) | 1399 (18.98) | 606 (21.29) | 567 (19.92) |
| **Psychopathology, count (%)** |  |  |  |  |  |  |
| AD | 458 (5.05) | 445 (4.9) | 269 (3.65) | 299 (4.06) | - | - |
| ADHD | 885 (9.75) | 450 (4.96) | 687 (9.32) | 313 (4.25) | - | - |
| CD | 174 (1.92) | 76 (0.84) | 38 (0.52) | 20 (0.27) | - | - |
| DD | 21 (0.23) | 14 (0.15) | 24 (0.33) | 26 (0.35) | - | - |
| OCD | 400 (4.41) | 269 (2.96) | 214 (2.9) | 199 (2.7) | - | - |
| ODD | 317 (3.49) | 189 (2.08) | 203 (2.75) | 143 (1.94) | - | - |

### Brain age models

Using TD children only (Figure 1A), we built eight tree-boosting models, one for each region depicted in Figure 1C, that included both male and female participants. All models performed well at predicting chronological age, returning MAE values ranging from 0.88 to 3.31 months and Pearson correlation coefficients ranging from 0.85 to 0.99 after age-bias correction (Table 2). Since males and females exhibit structural differences(63), we also built 16 additional models, one for each sex and brain region. Male-only models performed relatively similarly to the combined male/female models, with MAE values ranging from 0.93 to 3.35 months and Pearson correlation coefficients ranging from 0.86 to 0.99. In contrast, female-only models exhibited slightly higher MAE values, ranging from 1.28 to 5.25 months, and lower Pearson correlation coefficients, ranging from 0.73 to 0.98 (Supplementary Table 2), primarily driven by the temporal region model (MAE: 5.25 months, *r* = 0.73), which performed significantly worse than all other models.

**Table 2.** Model’s performance on the test set, before and after age-bias correction. ^1^: Statistics after correction as described by Beheshti *et al.* (2019). MAE: Mean absolute error (in months). r: Pearson’s correlation coefficient.

| Models | Before correction |  | After correction<sup>1</sup> |  |
| --- | --- | --- | --- | --- |
|  | MAE | r (p-value) | MAE | r (p-value) |
| Whole brain | 5.99 | 0.36 (< 0.001) | 3.31 | 0.85 (< 0.001) |
| Subcortical | 6.34 | 0.23 (< 0.001) | 1.78 | 0.96 (< 0.001) |
| Frontal | 6.21 | 0.28 (< 0.001) | 1.90 | 0.95 (< 0.001) |
| Temporal | 6.54 | 0.11 (< 0.001) | 1.08 | 0.99 (< 0.001) |
| Parietal | 6.48 | 0.17 (< 0.001) | 0.88 | 0.99 (< 0.001) |
| Insula | 6.53 | 0.11 (< 0.001) | 0.76 | 0.99 (< 0.001) |
| Limbic | 6.54 | 0.10 (< 0.001) | 3.00 | 0.95 (< 0.001) |
| Occipital | 6.42 | 0.19 (< 0.001) | 1.13 | 0.98 (< 0.001) |

**Table 3.** Clinical samples group effects for each brain region, controlled for sex, age, handedness, ethnicity, parental education, and family income. *d*: Cohen’s d effect size. AD: Anxiety disorder. ADHD: Attention-deficit/hyperactivity disorder. CD: Conduct disorder. OCD: Obsessive-compulsive disorder. ODD: Oppositional defiant disorder. DD: Depressive disorder. PSYPATHO: Participants with at least one psychiatric disorder. *p_fdr_*: FDR-corrected p-value (< 0.05 are **bolded**). All psychopathology diagnoses are derived from the KSADS and do not reflect a clinical diagnosis.

| Regions | AD |  | ADHD |  | CD |  | DD |  | OCD |  | ODD |  | PSYPATHO |  |
| --- | --- | --- | --- | --- | --- | --- | --- | --- | --- | --- | --- | --- | --- | --- |
| | $d$ | $p_{fdr}$ | $d$ | $p_{fdr}$ | $d$ | $p_{fdr}$ | $d$ | $p_{fdr}$ | $d$ | $p_{fdr}$ | $d$ | $p_{fdr}$ | $d$ | $p_{fdr}$ |
| Whole Brain | 0.03 | 0.657 | **-0.08** | **0.008** | -0.03 | 0.828 | -0.21 | 0.701 | -0.04 | 0.786 | -0.01 | 0.892 | -0.03 | 0.437 |
| Frontal | 0.00 | 0.979 | **-0.10** | **0.002** | -0.06 | 0.728 | -0.36 | 0.321 | -0.05 | 0.786 | -0.03 | 0.681 | -0.04 | 0.222 |
| Temporal | 0.03 | 0.657 | **-0.10** | **0.002** | 0.05 | 0.769 | 0.07 | 0.855 | 0.02 | 0.902 | 0.00 | 0.964 | -0.03 | 0.437 |
| Parietal | 0.05 | 0.595 | -0.03 | 0.263 | -0.01 | 0.928 | 0.03 | 0.855 | -0.01 | 0.995 | -0.09 | 0.241 | 0.00 | 0.825 |
| Occipital | 0.07 | 0.468 | -0.03 | 0.263 | 0.01 | 0.928 | -0.19 | 0.701 | -0.03 | 0.840 | -0.08 | 0.241 | 0.01 | 0.653 |
| Subcortical | 0.00 | 0.979 | **-0.12** | **0.002** | -0.17 | 0.083 | -0.06 | 0.855 | -0.10 | 0.120 | -0.09 | 0.241 | **-0.07** | **0.024** |
| Limbic | 0.05 | 0.595 | 0.00 | 0.440 | 0.15 | 0.094 | -0.17 | 0.701 | -0.01 | 0.995 | -0.04 | 0.681 | 0.00 | 0.825 |
| Insula | -0.02 | 0.676 | -0.02 | 0.263 | 0.10 | 0.346 | -0.06 | 0.855 | 0.00 | 0.995 | -0.05 | 0.540 | -0.02 | 0.474 |

### BAG associations with clinical diagnoses

To investigate BAG differences between clinical samples and TD children, we performed multivariate linear regression to extract group effect sizes while controlling for age, sex, handedness, ethnicity, parental education, and familial income. Using the BAG derived from the combined male/female model, participants suspected for ADHD showed lower whole-brain, frontal, temporal, and subcortical BAG compared to TD children (Cohen’s *d* = -0.08, *p_fdr_* = 0.008; *d* = -0.10, *p_fdr_*= 0.002; *d* = -0.10, *p_fdr_* = 0.002; *d* = -0.12, *p_fdr_* = 0.002, respectively) suggesting a delayed development in those regions. Participants suspected of at least one psychiatric disorder showed lower subcortical BAG (*d* = -0.07, *p* = 0.024), suggesting a delayed development of subcortical structures across all disorders. The remaining regions showed no significant differences between groups (Table 2). Sensitivity analyses were conducted using the BAG values derived from male-only and female-only models. Globally, most effect size trends were maintained in the sex-specific models but did not reach statistical significance after FDR correction (Supplementary Table 3).

### BAG associations with cognitive and behavioral profiles

We evaluated the relationship between BAG and baseline cognitive and behavioral profiles using multivariate linear regression models, controlling for age, sex, handedness, ethnicity, parental education, and family income (Table 4). Participants with higher membership values in the profile HC/LB (high cognition and low behavior) showed higher BAG values across multiple brain regions (whole brain, frontal, temporal, subcortical, limbic, and insula; *r_partial_* = 0.02-0.05), suggesting accelerated brain maturation. The opposite, i.e., delayed maturation, was found for the profile LC/LB (low cognition and low behavior) in all previously significant regions (*r_partial_* = -0.04 to -0.02), except for the insula and temporal regions. Participants with higher membership values in profile MC/HE (characterized by high externalizing behaviors and moderate cognition) exhibited lower BAG values in the frontal regions, suggesting delayed maturation of the frontal cortex (*r_partial_* = -0.03). Sensitivity analyses using BAG values from male and female-only brain age models yielded similar partial correlation coefficients but did not meet the statistical significance threshold after FDR correction. All statistics for the sensitivity analyses are presented in Supplementary Table 4.

**Table 4.** Partial correlation coefficient (*r_partial_*) between BAG values per regions derived from the combined male/female model and the membership values for each fuzzy cognitive and behavioral profile. MC/HSI: Moderate cognition and high stress/internalization. MC/HE: Moderate cognition and high externalization. HC/LB: High cognition and low behavior. LC/LB: Low cognition and low behavior. *p_fdr_*: FDR-corrected p-value (< 0.05 are **bolded**).

BAG associations with cognitive and behavioral profiles
| | $r_{partial}$ | $p_{fdr}$ | $r_{partial}$ | $p_{fdr}$ | $r_{partial}$ | $p_{fdr}$ | $r_{partial}$ | $p_{fdr}$ |
| --- | --- | --- | --- | --- | --- | --- | --- | --- |
| Whole Brain | -0.01 | 0.644 | -0.02 | 0.274 | **0.05** | **< 0.001** | **-0.04** | **< 0.001** |
| Frontal | -0.01 | 0.644 | **-0.03** | **0.016** | **0.05** | **< 0.001** | **-0.03** | **0.008** |
| Temporal | -0.01 | 0.557 | -0.02 | 0.291 | **0.02** | **0.034** | -0.02 | 0.057 |
| Parietal | -0.01 | 0.557 | -0.01 | 0.449 | -0.01 | 0.211 | 0.02 | 0.060 |
| Occipital | -0.03 | 0.086 | -0.02 | 0.291 | 0.02 | 0.071 | -0.01 | 0.322 |
| Subcortical | 0.00 | 0.861 | -0.01 | 0.301 | **0.04** | **< 0.001** | **-0.02** | **0.042** |
| Limbic | -0.01 | 0.557 | 0.00 | 0.775 | **0.03** | **0.002** | **-0.04** | **0.004** |
| Insula | 0.00 | 0.922 | -0.01 | 0.449 | **0.02** | **0.016** | -0.01 | 0.340 |

### BAG associations with longitudinal trajectories

We modeled membership values for each profile over time using a multivariate joint latent-class mixed model, extracting three latent trajectory classes that capture the evolution of profiles as a function of age (Figure 2). Trajectory class E (towards externalizing) showed a drastic movement from profile HC/LB to profile MC/HE, while trajectory class I (towards internalizing) represents similar movement amplitude, but from profile LC/LB to profile MC/HSI. The remaining trajectory class S (stable) showed little change with age, suggesting a more stable trajectory. Using the stable trajectory as the reference group, we evaluated how baseline BAG values relate to the chance of being assigned to either trajectory class E or class I. An increase in whole-brain, frontal, parietal, temporal, and insula BAG values significantly reduced the odds of being part of class E in male-only, female-only, and/or combined models (OR: 0.79 to 0.97, Figure 3). On the other hand, an increase in whole-brain, frontal, temporal, limbic, and subcortical BAG values increased the odds of following trajectory class I in male-only, female-only, and/or combined models (OR: 1.04 to 1.24, Figure 3). Overall, although not reaching statistical significance, an OR < 1 was observed for most BAG values in class E versus class S models, suggesting that accelerated brain maturation may be a protective factor for the development of externalization behaviors. Opposingly, a trend of OR > 1 was observed for BAG values in class I versus class S models, suggesting that accelerated brain maturation may be a contributing factor to the appearance of internalizing behaviors.

**Figure 2.**
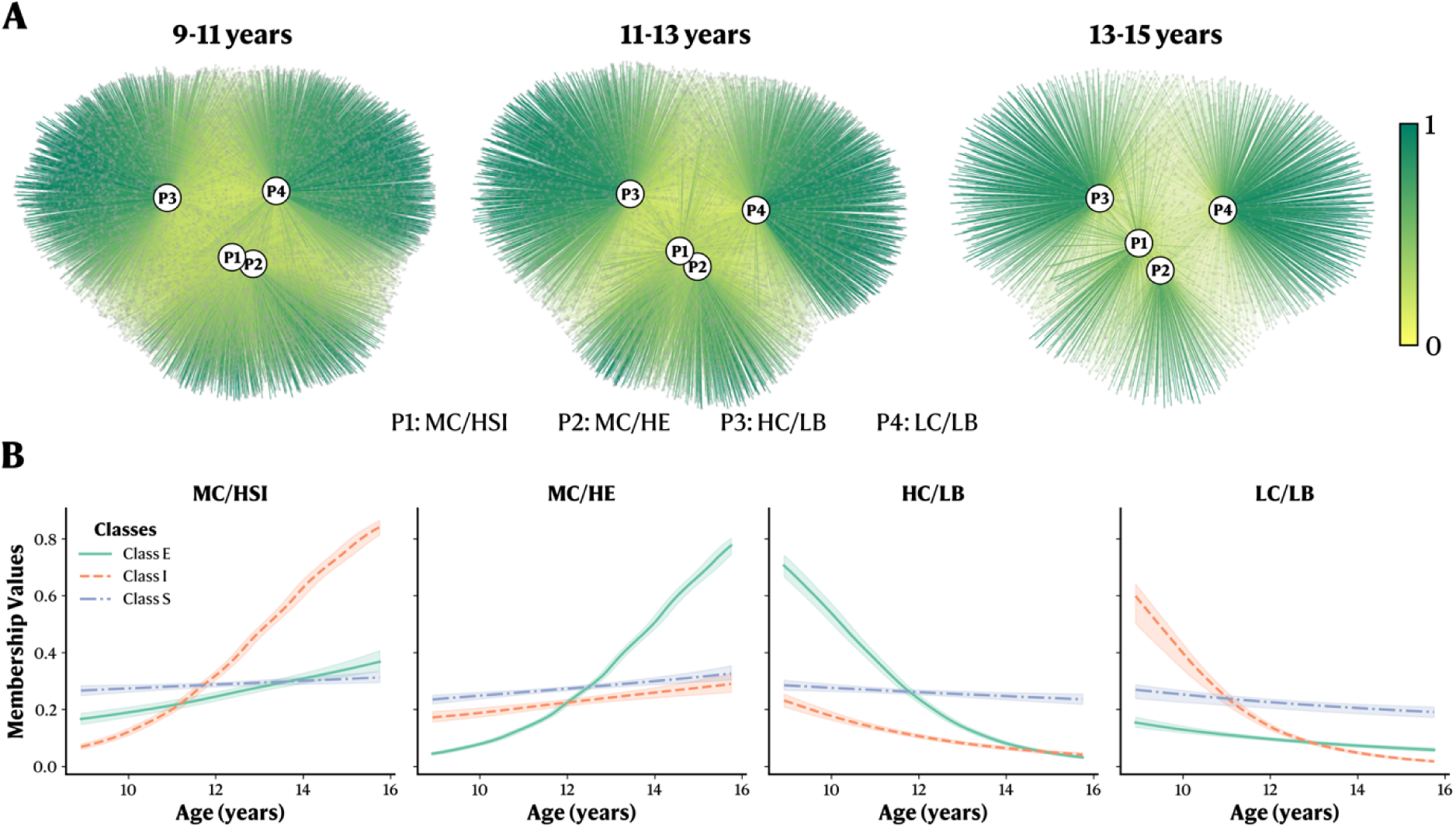
Cognitive and behavioral profiles for the 9-11, 11-13, and 13-15-year-old follow-ups, as well as the participants’ longitudinal trajectory between each time point. **A.** Graph network showing the distribution per time point of each profile. Graphs include all available participants, not just those retained for the longitudinal analysis. Gray nodes represent participants, while edges represent the membership value to a specific profile (darker green indicates a membership value closer to 1, while yellow edges indicate membership values close to 0). **B.** Latent trajectory classes derived from a multivariate joint latent class mixed model. Each plot represents the membership values of a profile based on participants’ age. Shaded areas surrounding the lines represent the 95% confidence interval.

**Figure 3.**
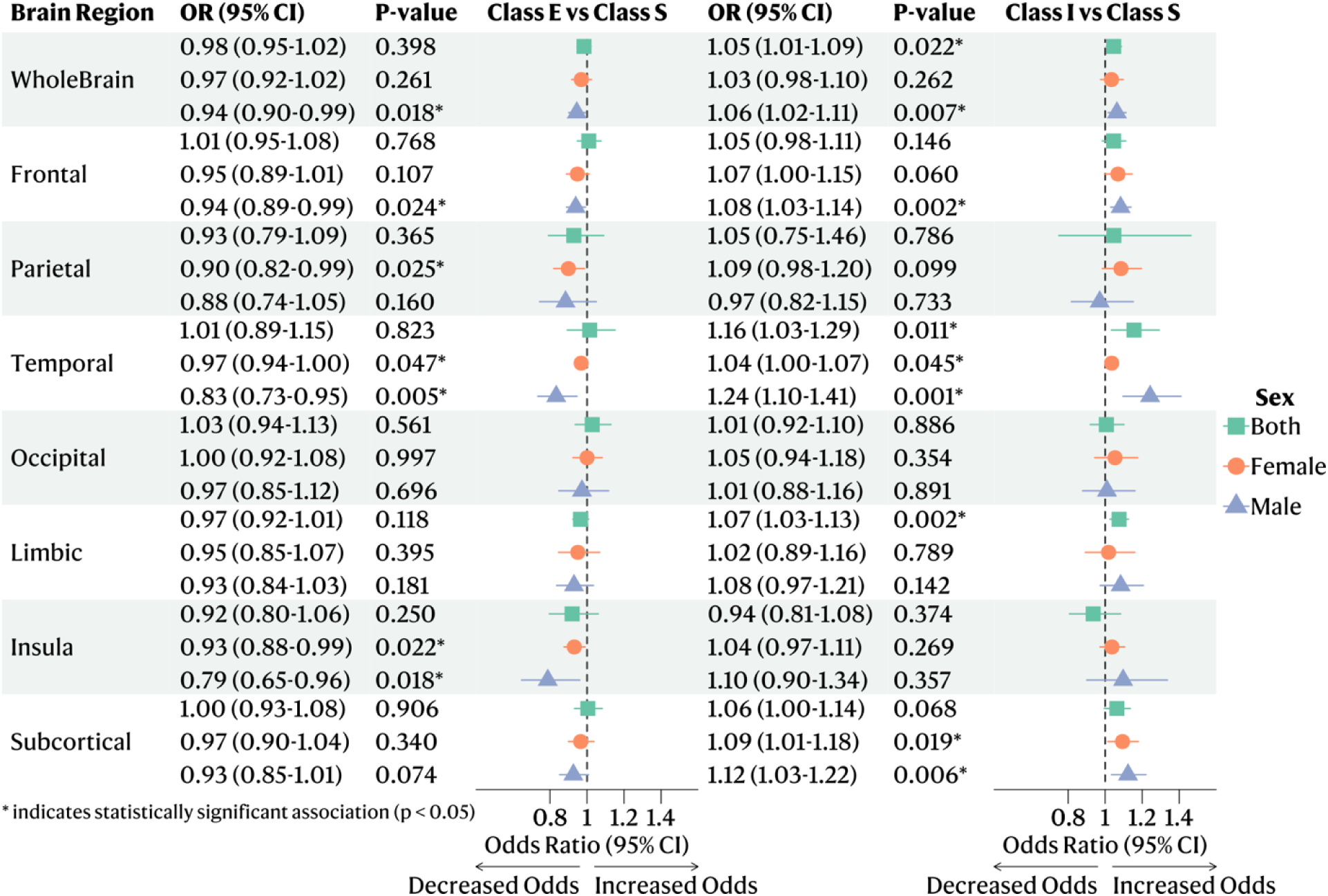
Forest plot of BAG Odds Ratio (OR) with 95% confidence interval of trajectory class E (towards externalizing behaviors) and trajectory class I (towards internalizing behaviors) compared to the reference stable class S. Results are stratified by sex. *: indicates statistically significant association. CI: Confidence Intervals.

## Discussion

In this large longitudinal pediatric cohort, we investigated the association between deviations in BAG and psychiatric diagnoses, continuous cognitive-behavioral profiles, and future developmental trajectories. Our results suggest that children suspected of ADHD showed delayed maturation in multiple brain regions, consistent with prior reports(13,53,64–66). Using novel continuous cognitive and behavioral profiles(27), we found that accelerated development was associated with a high cognition/low symptom profile, replicating results from multiple studies(18–20). In contrast, delayed development was observed in individuals in the low cognition and low symptom profile. Importantly, we evaluated whether baseline BAG values could hint towards future developmental trajectories. We extracted trajectories within the continuous cognitive and behavioral profiles, showing either stable (i.e., no drastic changes), increasing externalizing behaviors, or increasing internalizing behaviors, similar to previously derived latent growth trajectories in the same population(67). When investigating how baseline BAG relates to these trajectories, we demonstrate that accelerated brain maturation is protective against the development of externalizing behaviors, whereas it increases the odds of developing internalizing behaviors. The latter contrasts with our initial hypothesis that negative outcomes would be associated with delayed development. However, these results are in line with previous longitudinal studies, which, when compared with expected normative trajectories(1), showed accelerated development associated with lower externalizing(65,68–70) and higher internalizing(71–74) behaviors. This is also consistent with clinical evidence on interventional therapies in adults, showing that mentalization-based psychotherapy decreases acting-out behaviors over time(75–77). Interestingly, our analysis of children converges on a possible prognostic value of higher internalization behaviors in reducing externalizing behaviors and demonstrates its anatomical basis.

The regional and directional patterns of accelerated or delayed maturation observed in this study were consistent across psychiatric disorders, continuous cognitive/behavioral profiles, and trajectory analyses. These patterns likely reflect variation in the underlying cellular and molecular architecture of cortical and subcortical development in youth. Deviations from typical development found throughout the brain may reflect perturbations, either favorable or unfavorable, of the dopaminergic pathways and the expression of microglia, astrocytes, oligodendrocyte progenitor cells, pyramidal cells, or interneurons, all of which are predictors of cortical and subcortical changes during this developmental period(78,79). Additionally, these pathways/cell types have been previously associated with ADHD(80–83), externalizing behaviors(84,85), and/or internalizing behaviors(84,85) in animal and human studies. Regional findings, particularly in frontal, temporal, or subcortical regions, highlighted areas enriched in these neurobiological markers(78). Considering the current findings, an accelerated maturation of these regions might reflect differential expression patterns that could facilitate the development of adaptive behavior and cognition, preventing trajectories towards externalizing behaviors while predisposing others to develop internalizing symptoms. While the current study cannot establish a causal, mechanistic link between brain maturation status and future developmental trajectories, it suggests that the BAG framework, through its macroscale evaluation of normative brain development, may index regionally specific molecular and cellular trajectories that control the timing and outcome of neurodevelopmental pathways.

A crucial critique of the BAG framework is its reliance on prediction errors from trained models. This prediction error can be affected by multiple sources, including the actual physiology, but also noise, which can originate from site effects and data quality(44). However, we mitigated this risk of prediction bias by using state-of-the-art harmonization methods and rigorous quality control (Supplementary Figures 2 to 11). Another concern with the BAG framework is its dependence on age, which systematically overestimates younger participants’ age while underestimating older participants’ age(86). This dependence was mitigated using correction techniques in the present study (Supplementary Figure 12). While those techniques have been shown to artificially inflate model performance(86), they have been found not to affect downstream effect sizes(47), assuming the inclusion of age in statistical models, as done in the present study. Our brain age models were trained on a narrow age range (∼9-11 years), which constrains the BAG relative to those trained on a broader age range. This might have led to our effect sizes being underestimated compared to those in other studies. We assessed case-control differences where participants were not diagnosed by clinicians, but rather through a parental questionnaire. While it is considered the gold standard in research settings(53), it might also have influenced our observed effect sizes. Additionally, while ABCD is designed to be representative of the US population(30), our study was conducted with participants with a North American background; future studies should focus on replication in other populations of diverse ethnicities.

From a clinical perspective, our findings suggest that the BAG framework may be useful in the development of precision psychiatry. One contribution could be in prognosis, since either delayed or accelerated development is associated with two different trajectories towards psychopathology. Precision psychiatry aims to predict trajectories efficiently to widen the window for potential interventions(16). BAG could help identify patients with a predisposition to the development of internalization symptoms (or externalization symptoms), thereby prompting clinicians to consider preventive interventions or closer monitoring. While the BAG framework holds promise for precision psychiatry, widespread adoption would require additional studies to validate its performance across different populations (e.g., to avoid cultural biases) and in the presence of noisy, missing, or sparse data(16,17), which is often the case in clinical contexts. Nonetheless, brain age models could represent a viable approach to improve precision medicine in psychiatry.

In summary, we applied the brain age framework to a large pediatric population. We investigated brain age differences in psychiatric disorders and their relationships with state-of-the-art continuous cognitive and behavioral profiles and developmental trajectories. We found delayed development in youth with ADHD in multiple regions, and accelerated maturation in participants with higher cognitive abilities. Accelerated maturation was also predictive of a trajectory towards internalizing behaviors, while also acting as a protective factor for the development of externalizing behaviors. Clinically, these results highlight the potential of brain age models to serve as an individualized developmental biomarker, capable of informing early identification of at-risk youth. Incorporating brain age within precision psychiatry may help clinicians anticipate divergent developmental trajectories, widening the window for timely interventions.

## Data and Code Availability

The Adolescent Brain and Cognitive Development (ABCD) and the Boston Adolescent Neuroimaging of Depression and Anxiety (BANDA) are freely available datasets accessible via the NIH Brain Development Cohorts (NBDC) Data Sharing Platform (https://www.nbdc-datahub.org/). Researchers must apply for a Data Use Certificate, which will be reviewed and granted by study administrators. Code to reproduce the findings, including data curation, preprocessing, analysis, and visualization, is accessible at https://github.com/gagnonanthony/Gagnon_BrainAge_2025.

## Acknowledgments.

Data used in the preparation of this article were obtained from the Adolescent Brain Cognitive Development^SM^ (ABCD) Study (https://abcdstudy.org), held in the NIMH Data Archive (NDA). This is a multisite, longitudinal study designed to recruit more than 10,000 children age 9-10 and follow them over 10 years into early adulthood. The ABCD Study® is supported by the National Institutes of Health and additional federal partners under award numbers U01DA041048, U01DA050989, U01DA051016, U01DA041022, U01DA051018, U01DA051037, U01DA050987, U01DA041174, U01DA041106, U01DA041117, U01DA041028, U01DA041134, U01DA050988, U01DA051039, U01DA041156, U01DA041025, U01DA041120, U01DA051038, U01DA041148, U01DA041093, U01DA041089, U24DA041123, U24DA041147. A full list of supporters is available at https://abcdstudy.org/federal-partners.html. A listing of participating sites and a complete listing of the study investigators can be found at https://abcdstudy.org/consortium_members/. ABCD consortium investigators designed and implemented the study and/or provided data but did not necessarily participate in the analysis or writing of this report. This manuscript reflects the views of the authors and may not reflect the opinions or views of the NIH or ABCD consortium investigators. The ABCD data repository grows and changes over time. The ABCD data used in this report came from <u>10.15154/z563-zd24</u>. DOIs can be found at https://nda.nih.gov/abcd/abcd-annual-releases. AG was supported by a Canadian Institute of Health Research Doctoral Award (#493956). MAB is supported by a Junior 1 career award from the Fonds de Recherche du Québec – Santé (FRQS).

## Author Contributions

AG, MAB, MD, and LT contributed to the study design and conceptualization. AG conducted the analyses and wrote the final draft. All authors contributed to the interpretation of findings and approved the final version of the manuscript before submission. AG and LT were responsible for the decision to publish.

## References

1. Bethlehem RAI, Seidlitz J, White SR, Vogel JW, Anderson KM, Adamson C, et al. (2022): Brain charts for the human lifespan. Nature 604: 525–533.

2. Silk TJ, Wood AG (2011): Lessons About Neurodevelopment From Anatomical Magnetic Resonance Imaging. Journal of Developmental & Behavioral Pediatrics 32: 158–168.

3. Schumann G, Benegal V, Yu C, Tao S, Jernigan T, Heinz A, et al. (2019): Precision medicine and global mental health. The Lancet Global Health 7: e32.

4. Jones PB (2013): Adult mental health disorders and their age at onset. Br J Psychiatry 202: s5–s10.

5. Marín O (2016): Developmental timing and critical windows for the treatment of psychiatric disorders. Nat Med 22: 1229–1238.

6. Prince M, Patel V, Saxena S, Maj M, Maselko J, Phillips MR, Rahman A (2007): No health without mental health. The Lancet 370: 859–877.

7. Dosenbach NUF, Nardos B, Cohen AL, Fair DA, Power JD, Church JA, et al. (2010): Prediction of Individual Brain Maturity Using fMRI. Science 329: 1358–1361.

8. Franke K, Ziegler G, Klöppel S, Gaser C (2010): Estimating the age of healthy subjects from T1-weighted MRI scans using kernel methods: Exploring the influence of various parameters. NeuroImage 50: 883–892.

9. Cole JH, Franke K (2017): Predicting Age Using Neuroimaging: Innovative Brain Ageing Biomarkers. Trends in Neurosciences 40: 681–690.

10. Chung Y, Addington J, Bearden CE, Cadenhead K, Cornblatt B, Mathalon DH, et al. (2018): Use of Machine Learning to Determine Deviance in Neuroanatomical Maturity Associated With Future Psychosis in Youths at Clinically High Risk. JAMA Psychiatry 75: 960–968.

11. Tunç B, Yankowitz LD, Parker D, Alappatt JA, Pandey J, Schultz RT, Verma R (2019): Deviation from normative brain development is associated with symptom severity in autism spectrum disorder. Molecular Autism 10: 46.

12. Xie Y, Sun J, Man W, Zhang Z, Zhang N (2023): Personalized estimates of brain cortical structural variability in individuals with Autism spectrum disorder: the predictor of brain age and neurobiology relevance. Molecular Autism 14: 27.

13. Kozhemiako N, Buckley AW, Chervin RD, Redline S, Purcell SM (2024): Mapping neurodevelopment with sleep macro- and micro-architecture across multiple pediatric populations. NeuroImage: Clinical 41: 103552.

14. Lund MJ, Alnæs D, De Lange A-MG, Andreassen OA, Westlye LT, Kaufmann T (2022): Brain age prediction using fMRI network coupling in youths and associations with psychiatric symptoms. NeuroImage: Clinical 33: 102921.

15. Luna A, Bernanke J, Kim K, Aw N, Dworkin JD, Cha J, Posner J (2021): Maturity of gray matter structures and white matter connectomes, and their relationship with psychiatric symptoms in youth. Human Brain Mapping 42: 4568–4579.

16. Hauser TU, Skvortsova V, De Choudhury M, Koutsouleris N (2022): The promise of a model-based psychiatry: building computational models of mental ill health. The Lancet Digital Health 4: e816–e828.

17. Grzenda A, Kraguljac NV, McDonald WM, Nemeroff C, Torous J, Alpert JE, et al. (2021): Evaluating the Machine Learning Literature: A Primer and User’s Guide for Psychiatrists. AJP 178: 715–729.

18. Erus G, Battapady H, Satterthwaite TD, Hakonarson H, Gur RE, Davatzikos C, Gur RC (2015): Imaging Patterns of Brain Development and their Relationship to Cognition. Cerebral Cortex 25: 1676–1684.

19. Ullman H, Klingberg T (2016): Timing of White Matter Development Determines Cognitive Abilities at School Entry but Not in Late Adolescence. Cereb Cortex cercor;bhw256v2.

20. Lewis JD, Evans AC, Tohka J (2018): T1 white/gray contrast as a predictor of chronological age, and an index of cognitive performance. NeuroImage 173: 341–350.

21. Ng C, Huang P, Cho Y, Lee P, Liu Y, Chang T (2024): Frontoparietal and salience network synchronizations during nonsymbolic magnitude processing predict brain age and mathematical performance in youth. Human Brain Mapping 45: e26777.

22. Ball G, Kelly CE, Beare R, Seal ML (2021): Individual variation underlying brain age estimates in typical development. NeuroImage 235: 118036.

23. Ball G, Adamson C, Beare R, Seal ML (2017): Modelling neuroanatomical variation during childhood and adolescence with neighbourhood-preserving embedding. Sci Rep 7: 17796.

24. Thompson WK, Barch DM, Bjork JM, Gonzalez R, Nagel BJ, Nixon SJ, Luciana M (2019): The structure of cognition in 9 and 10 year-old children and associations with problem behaviors: Findings from the ABCD study’s baseline neurocognitive battery. Developmental Cognitive Neuroscience 36: 100606.

25. Moore DM, Conway ARA (2023): The Structure of Cognitive Abilities and Associations with Problem Behaviors in Early Adolescence: An Analysis of Baseline Data from the Adolescent Brain Cognitive Development Study. J Intell 11: 90.

26. Pines A, Tozzi L, Bertrand C, Keller AS, Zhang X, Whitfield-Gabrieli S, et al. (2024): Psychiatric Symptoms, Cognition, and Symptom Severity in Children. JAMA Psychiatry. 10.1001/jamapsychiatry.2024.2399

27. Gagnon A, Gillet V, Desautels A-S, Lepage J-F, Baccarelli AA, Posner J, et al. (2025): Beyond discrete classifications: a computational approach to the continuum of cognition and behavior in children. npj Mental Health Res 4: 48.

28. Cuthbert BN, Insel TR (2013): Toward the future of psychiatric diagnosis: the seven pillars of RDoC. BMC Med 11: 126.

29. Kotov R, Krueger RF, Watson D, Achenbach TM, Althoff RR, Bagby RM, et al. (2017): The Hierarchical Taxonomy of Psychopathology (HiTOP): A dimensional alternative to traditional nosologies. Journal of Abnormal Psychology 126: 454–477.

30. Garavan H, Bartsch H, Conway K, Decastro A, Goldstein RZ, Heeringa S, et al. (2018): Recruiting the ABCD sample: Design considerations and procedures. Developmental Cognitive Neuroscience 32: 16–22.

31. Casey BJ, Cannonier T, Conley MI, Cohen AO, Barch DM, Heitzeg MM, et al. (2018): The Adolescent Brain Cognitive Development (ABCD) study: Imaging acquisition across 21 sites. Developmental Cognitive Neuroscience 32: 43–54.

32. Li W, Fan L, Shi W, Lu Y, Li J, Luo N, et al. (2023): Brainnetome atlas of preadolescent children based on anatomical connectivity profiles. Cerebral Cortex 33: 5264–5275.

33. Cetin-Karayumak S, Zhang F, Zurrin R, Billah T, Zekelman L, Makris N, et al. (2024): Harmonized diffusion MRI data and white matter measures from the Adolescent Brain Cognitive Development Study. Sci Data 11: 249.

34. Henschel L, Conjeti S, Estrada S, Diers K, Fischl B, Reuter M (2020): FastSurfer - A fast and accurate deep learning based neuroimaging pipeline. NeuroImage 219: 117012.

35. Di Tommaso P, Chatzou M, Floden EW, Barja PP, Palumbo E, Notredame C (2017): Nextflow enables reproducible computational workflows. Nat Biotechnol 35: 316–319.

36. Ewels PA, Peltzer A, Fillinger S, Patel H, Alneberg J, Wilm A, et al. (2020): The nf-core framework for community-curated bioinformatics pipelines. Nature Biotechnology 38: 276–278.

37. Ewels P, Magnusson M, Lundin S, Käller M (2016): MultiQC: summarize analysis results for multiple tools and samples in a single report. Bioinformatics 32: 3047–3048.

38. Kurtzer GM, Sochat V, Bauer MW (2017): Singularity: Scientific containers for mobility of compute ((A. Gursoy, editor)). PLoS ONE 12: e0177459.

39. Klein A, Ghosh SS, Avants B, Yeo BTT, Fischl B, Ardekani B, et al. (2010): Evaluation of volume-based and surface-based brain image registration methods. NeuroImage 51: 214–220.

40. Fischl B (2012): FreeSurfer. NeuroImage 62: 774–781.

41. Fortin J-P, Cullen N, Sheline YI, Taylor WD, Aselcioglu I, Cook PA, et al. (2018): Harmonization of cortical thickness measurements across scanners and sites. NeuroImage 167: 104–120.

42. Girard G, Edde M, Dumais F, David Y, Dumont M, Theaud G, et al. (2025, November 6): Clinical-ComBAT: a diffusion-weighted MRI harmonization method for clinical applications [no. arXiv:2511.04871]. arXiv. 10.48550/arXiv.2511.04871

43. Chen T, Guestrin C (2016): XGBoost: A Scalable Tree Boosting System. Proceedings of the 22nd ACM SIGKDD International Conference on Knowledge Discovery and Data Mining 785–794.

44. Kaufmann T, van der Meer D, Doan NT, Schwarz E, Lund MJ, Agartz I, et al. (2019): Common brain disorders are associated with heritable patterns of apparent aging of the brain. Nature Neuroscience 22: 1617–1623.

45. Beck D, Whitmore L, MacSweeney N, Brieant A, Karl V, De Lange A-MG, et al. (2025): Dimensions of Early-Life Adversity Are Differentially Associated With Patterns of Delayed and Accelerated Brain Maturation. Biological Psychiatry 97: 64–72.

46. Dehestani N, Whittle S, Vijayakumar N, Silk TJ (2023): Developmental brain changes during puberty and associations with mental health problems. Developmental Cognitive Neuroscience 60: 101227.

47. De Lange A-MG, Cole JH (2020): Commentary: Correction procedures in brain-age prediction. NeuroImage: Clinical 26: 102229.

48. Beheshti I, Nugent S, Potvin O, Duchesne S (2019): Bias-adjustment in neuroimaging-based brain age frameworks: A robust scheme. NeuroImage: Clinical 24: 102063.

49. Le TT, Kuplicki RT, McKinney BA, Yeh H-W, Thompson WK, Paulus MP, Tulsa 1000 Investigators (2018): A Nonlinear Simulation Framework Supports Adjusting for Age When Analyzing BrainAGE. Front Aging Neurosci 10: 317.

50. Smith SM, Vidaurre D, Alfaro-Almagro F, Nichols TE, Miller KL (2019): Estimation of brain age delta from brain imaging. NeuroImage 200: 528–539.

51. Niu X, Zhang F, Kounios J, Liang H (2020): Improved prediction of brain age using multimodal neuroimaging data. Human Brain Mapping 41: 1626–1643.

52. Liang H, Zhang F, Niu X (2019): Investigating systematic bias in brain age estimation with application to post-traumatic stress disorders. Human Brain Mapping 40: 3143–3152.

53. Bernanke J, Luna A, Chang L, Bruno E, Dworkin J, Posner J (2022): Structural brain measures among children with and without ADHD in the Adolescent Brain and Cognitive Development Study cohort: a cross-sectional US population-based study. The Lancet Psychiatry 9: 222–231.

54. Gershon RC, Wagster MV, Hendrie HC, Fox NA, Cook KF, Cindy J. Nowinski (2013): NIH toolbox for assessment of neurological and behavioral function. Neurology 80: S2–S6.

55. Acker WL, Acker C, England NF for ER in, Wales, Psychiatry U of LI of (1982): Bexley Maudsley Automated Psychological Screening and Bexley Maudsley Category Sorting Test Manual. NFER-Nelson, for the Institute of Psychiatry. Retrieved from https://books.google.ca/books?id=h-XaZwEACAAJ

56. Luciana M, Bjork JM, Nagel BJ, Barch DM, Gonzalez R, Nixon SJ, Banich MT (2018): Adolescent neurocognitive development and impacts of substance use: Overview of the adolescent brain cognitive development (ABCD) baseline neurocognition battery. Developmental Cognitive Neuroscience 32: 67–79.

57. Wechsler D (2014): Wechsler intelligence scale for children–Fifth Edition (WISC-V). Bloomington, MN: Pearson.

58. Achenbach TM, Edelbrock C (1991): Child behavior checklist. Burlington (Vt) 7: 371–392.

59. Nakagawa S, Cuthill IC (2007): Effect size, confidence interval and statistical significance: a practical guide for biologists. Biological Reviews 82: 591–605.

60. Good P (2000): Permutation Tests. New York, NY: Springer. 10.1007/978-1-4757-3235-1

61. Benjamini Y, Krieger AM, Yekutieli D (2006): Adaptive linear step-up procedures that control the false discovery rate. Biometrika 93: 491–507.

62. Proust-Lima C, Philipps V, Liquet B (2017): Estimation of Extended Mixed Models Using Latent Classes and Latent Processes: The R Package lcmm. Journal of Statistical Software 78: 1–56.

63. Ritchie SJ, Cox SR, Shen X, Lombardo MV, Reus LM, Alloza C, et al. (2018): Sex Differences in the Adult Human Brain: Evidence from 5216 UK Biobank Participants. Cerebral Cortex 28: 2959–2975.

64. Shaw P, Eckstrand K, Sharp W, Blumenthal J, Lerch JP, Greenstein D, et al. (2007): Attention-deficit/hyperactivity disorder is characterized by a delay in cortical maturation. Proceedings of the National Academy of Sciences 104: 19649–19654.

65. Townend S, Staginnus M, Gao Y, Alexander N, Arolt V, Banaschewski T, et al. (2025): Shared and distinct alterations in brain structure of youth with internalizing or externalizing disorders: Findings from the ENIGMA Antisocial Behavior, ADHD, MDD, and Anxiety Working Groups. Biological Psychiatry. 10.1016/j.biopsych.2025.08.003

66. Kurth F, Levitt JG, Gaser C, Alger J, Loo SK, Narr KL, et al. (2022): Preliminary evidence for a lower brain age in children with attention-deficit/hyperactivity disorder. Front Psychiatry 13: 1019546.

67. Brieant A, Cai T, Ip KI, Holt-Gosselin B, Gee DG (2025): Heterogeneity in Developmental Trajectories of Internalizing and Externalizing Symptomatology: Associations with Risk and Protective Factors. Child Psychiatry Hum Dev. 10.1007/s10578-024-01804-0

68. Bos MGN, Wierenga LM, Blankenstein NE, Schreuders E, Tamnes CK, Crone EA (2018): Longitudinal structural brain development and externalizing behavior in adolescence. Journal of Child Psychology and Psychiatry 59: 1061–1072.

69. Parkes L, Moore TM, Calkins ME, Cook PA, Cieslak M, Roalf DR, et al. (2021): Transdiagnostic dimensions of psychopathology explain individuals’ unique deviations from normative neurodevelopment in brain structure. Transl Psychiatry 11: 232.

70. Jarvers I, Kandsperger S, Schleicher D, Ando A, Resch F, Koenig J, et al. (2022): The relationship between adolescents’ externalizing and internalizing symptoms and brain development over a period of three years. NeuroImage: Clinical 36: 103195.

71. Zhao Y, Paulus Mp, Potenza Mn (2023): Brain structural co-development is associated with internalizing symptoms two years later in the ABCD cohort. Journal of Behavioral Addictions 12: 80–93.

72. MacSweeney N, Beck D, Whitmore L, Mills KL, Westlye LT, Soest T von, et al. (2024): Multimodal brain age indicators of internalising problems in early adolescence: A longitudinal investigation. Biological Psychiatry: Cognitive Neuroscience and Neuroimaging. 10.1016/j.bpsc.2024.11.003

73. Bos MGN, Peters S, Van De Kamp FC, Crone EA, Tamnes CK (2018): Emerging depression in adolescence coincides with accelerated frontal cortical thinning. Child Psychology Psychiatry 59: 994–1002.

74. Drobinin V, Van Gestel H, Helmick CA, Schmidt MH, Bowen CV, Uher R (2022): The Developmental Brain Age Is Associated With Adversity, Depression, and Functional Outcomes Among Adolescents. Biological Psychiatry: Cognitive Neuroscience and Neuroimaging 7: 406–414.

75. Fonagy P, Simes E, Yirmiya K, Wason J, Barrett B, Frater A, et al. (2025): Mentalisation-based treatment for antisocial personality disorder in males convicted of an offence on community probation in England and Wales (Mentalization for Offending Adult Males, MOAM): a multicentre, assessor-blinded, randomised controlled trial. The Lancet Psychiatry 12: 208–219.

76. Bateman A, O’Connell J, Lorenzini N, Gardner T, Fonagy P (2016): A randomised controlled trial of mentalization-based treatment versus structured clinical management for patients with comorbid borderline personality disorder and antisocial personality disorder. BMC Psychiatry 16: 304.

77. Anthony Bateman MA, Peter Fonagy PD (2009): Randomized Controlled Trial of Outpatient Mentalization-Based Treatment Versus Structured Clinical Management for Borderline Personality Disorder. American Journal of Psychiatry. 10.1176/appi.ajp.2009.09040539

78. Lotter LD, Saberi A, Hansen JY, Misic B, Paquola C, Barker GJ, et al. (2024): Regional patterns of human cortex development correlate with underlying neurobiology. Nat Commun 15: 7987.

79. Vidal-Pineiro D, Parker N, Shin J, French L, Grydeland H, Jackowski AP, et al. (2020): Cellular correlates of cortical thinning throughout the lifespan. Sci Rep 10: 21803.

80. Posner J, Polanczyk GV, Sonuga-Barke E (2020): Attention-deficit hyperactivity disorder. The Lancet 395: 450–462.

81. Bordeleau M, Carrier M, Luheshi GN, Tremblay M-È (2019): Microglia along sex lines: From brain colonization, maturation and function, to implication in neurodevelopmental disorders. Seminars in Cell & Developmental Biology 94: 152–163.

82. Hess JL, Radonjić NV, Patak J, Glatt SJ, Faraone SV (2021): Autophagy, apoptosis, and neurodevelopmental genes might underlie selective brain region vulnerability in attention-deficit/hyperactivity disorder. Molecular Psychiatry 26: 6643–6654.

83. Ferranti AS, Luessen DJ, Niswender CM (2024): Novel pharmacological targets for GABAergic dysfunction in ADHD. Neuropharmacology 249: 109897.

84. Hansen JY, Markello RD, Vogel JW, Seidlitz J, Bzdok D, Misic B (2021): Mapping gene transcription and neurocognition across human neocortex. Nat Hum Behav 5: 1240–1250.

85. Schmidt LA, Fox NA, Hamer DH (2007): Evidence for a gene–gene interaction in predicting children’s behavior problems: Association of serotonin transporter short and dopamine receptor D4 long genotypes with internalizing and externalizing behaviors in typically developing 7-year-olds. Development and Psychopathology 19: 1105–1116.

86. Butler ER, Chen A, Ramadan R, Le TT, Ruparel K, Moore TM, et al. (2021): Pitfalls in brain age analyses. Human Brain Mapping 42: 4092–4101.

