## Supplement for "Regional brain age deviations reveal divergent developmental pathways in youth"

**Gagnon A, *et al.***

**Table of contents**

**Supplementary Tables**

| **Supplementary Table 1.** Included parcels in each coarse area’s brain age model. | p. 3-5 |
| --- | --- |
| **Supplementary Table 2.** Male and female models’ performance on their respective test sets, before and after age-bias correction. | p. 6 |
| **Supplementary Table 3.** Clinical samples group effects for each brain region stratified by BAG derived from the male and female-only models | p. 7 |
| **Supplementary Table 4.** Partial correlation coefficient (*r_partial_*) between BAG values per regions derived from the male and female only models and the membership values for each fuzzy cognitive and behavioral profile. | p. 8 |

**Supplementary Figures**

| **Supplementary Figure 1.** Flowchart of participants’ exclusion. | p. 9 |
| --- | --- |
| **Supplementary Figure 2.** Distribution of subcortical volumes for left and right hemispheres before outlier removal. | p. 10 |
| **Supplementary Figure 3.** Distribution of subcortical volumes for left and right hemispheres after outlier removal. | p. 11 |
| **Supplementary Figure 4.** Distribution of cortical thickness, volume, and surface area for the **left hemisphere** before outlier removal. | p. 12 |
| **Supplementary Figure 5.** Distribution of cortical thickness, volume, and surface area for the **left hemisphere** after outlier removal. | p. 13 |
| **Supplementary Figure 6.** Distribution of cortical thickness, volume, and surface area for the **right hemisphere** before outlier removal. | p. 14 |
| **Supplementary Figure 7.** Distribution of cortical thickness, volume, and surface area for the **right hemisphere** after outlier removal. | p. 15 |
| **Supplementary Figure 8.** Overview of the participants’ site distribution, in addition to the characteristics of the reference site used as a target for data harmonization (site 16). | p. 16 |
| **Supplementary Figure 9.** Example visualization of the pairwise harmonization to the reference site (site 16) for the volume metric in the inferior parietal lobule A39rd region. | p. 17 |
| **Supplementary Figure 10.**  Example visualization of the pairwise harmonization to the reference site (site 16) for the cortical thickness metric in the inferior parietal lobule A39rd region. | p. 18 |
| **Supplementary Figure 11.**  Example visualization of the pairwise harmonization to the reference site (site 16) for the surface area metric in the inferior parietal lobule A39rd region. | p. 19 |
| **Supplementary Figure 12.** Visualization of the age bias in brain age prediction for the whole-brain model before and after correction. | p. 20 |
| **Supplementary Figure 13.** Statistical criteria plots assessing the fit of multiple latent class models (2 to 10). | p. 21 |

| **Coarse areas (metric(s) included)** | **Included parcels** |
| --- | --- |
| Subcortical/cerebellar (volume) | - Amyg_L(R)_2_1: mAmyg, medial amygdala - Amyg_L(R)_2_2: lAmyg, lateral amygdala - Hipp_L(R)_2_1: rHipp, rostral hippocampus - Hipp_L(R)_2_2: cHipp, caudal hippocampus - BG_L(R)_6_1: vCa, ventral caudate - BG_L(R)_6_2: GP, globus pallidum - BG_L(R)_6_3: NAC, nucleus accumbens - BG_L(R)_6_4: vmPu, ventromedial putamen - BG_L(R)_6_5: dCa, dorsal caudate - BG_L(R)_6_6: dlPu, dorsolateral putamen - Tha_L(R)_8_1: mPFtha, medial pre-frontal thalamus - Tha_L(R)_8_2: mPMtha, pre-motor thalamus - Tha_L(R)_8_3: Stha, sensory thalamus - Tha_L(R)_8_4: rTtha, rostral temporal thalamus - Tha_L(R)_8_5: PPtha, posterior parietal thalamus - Tha_L(R)_8_6: Otha, occipital thalamus - Tha_L(R)_8_7: cTtha, caudal temporal thalamus - Tha_L(R)_8_8: lPFtha, lateral pre-frontal thalamus - Right cerebellum cortex - Left cerebellum cortex - BrainStem |
| Frontal (volume, thickness, surface area) | - SFG_L(R)_6_1: A8m, medial area 8 - SFG_L(R)_6_2: A8dl, dorsolateral area 8 - SFG_L(R)_6_3: A9m/A9l, medial area 9/lateral area 9 - SFG_L(R)_6_4: A6dl, dorsolateral area 6 - SFG_L(R)_6_5: A6m, medial area 6 - SFG_L(R)_6_6: A10m, medial area 10 - MFG_L(R)_7_1: A9/46d, dorsal area 9/46 - MFG_L(R)_7_2: IFJ, inferior frontal junction - MFG_L(R)_7_3: A46, area 46 - MFG_L(R)_7_4: A9/46v, ventral area 9/46 - MFG_L(R)_7_5: A8vl, ventrolateral area 8 - MFG_L(R)_7_6: A6vl, ventrolateral area 6 - MFG_L(R)_7_7: A10l, lateral area 10 - IFG_L(R)_6_1: A44d, dorsal area 44 - IFG_L(R)_6_2: IFS, inferior frontal sulcus - IFG_L(R)_6_3: A45c, caudal area 45 - IFG_L(R)_6_4: A45r, rostral area 45 - IFG_L(R)_6_5: A44op, opercular area 44 - IFG_L(R)_6_6: A44v, ventral area 44 - OrG_L(R)_6_1: A14m, medial area 14 - OrG_L(R)_6_2: A12/47o, orbital area 12/47 - OrG_L(R)_6_3: A11l, lateral area 11 - OrG_L(R)_6_4: A11m, medial area 11 - OrG_L(R)_6_5: A13, area 13 - OrG_L(R)_6_6: A12/47l, lateral area 12/47 - PrG_L(R)_6_1: A4hf, area 4(head and face region) - PrG_L(R)_6_2: A6cdl, caudal dorsolateral area 6 - PrG_L(R)_6_3: A4ul, area 4(upper limb region) - PrG_L(R)_6_4: A4t, area 4(trunk region) - PrG_L(R)_6_5: A4tl, area 4(tongue and larynx region) - PrG_L(R)_6_6: A6cvl, caudal ventrolateral area 6 - PCL_L(R)_2_1: A1/2/3ll, area 1/2/3(lower limb region) - PCL_L(R)_2_2: A4ll, area 4 |
| Temporal (volume, thickness, surface area) | - STG_L(R)_6_1: A38m, medial area 38 - STG_L(R)_6_2: A41/42, area 41/42 - STG_L(R)_6_3: TE1.0 and TE1.2 - STG_L(R)_6_4: A22c, caudal area 22 - STG_L(R)_6_5: A38l, lateral area 38 - STG_L(R)_6_6: A22r, rostral area 22 - MTG_L(R)_4_1: A21c, caudal area 21 - MTG_L(R)_4_2: A21r, rostral area 21 - MTG_L(R)_4_3: A37dl, dorsolateral area 37 - MTG_L(R)_4_4: aSTS, anterior superior temporal sulcus - ITG_L(R)_5_1: A37elv, extreme lateroventral area 37 - ITG_L(R)_5_2: A20r, rostral area 20 - ITG_L(R)_5_3: A20il, intermediate lateral area 20 - ITG_L(R)_5_4: A20cl, caudolateral of area 20 - ITG_L(R)_5_5: A20cv, caudoventral of area 20 - FuG_L(R)_3_1: A20rv, rostroventral area 20 - FuG_L(R)_3_2: A37mv, medioventral area 37 - FuG_L(R)_3_3: A37lv, lateroventral area 37 - PhG_L(R)_6_1: A35/36r, rostral area 35/36 - PhG_L(R)_6_2: A35/36c, caudal area 35/36 - PhG_L(R)_6_3: TL, area TL - PhG_L(R)_6_4: A28/34, area 28/34 - PhG_L(R)_6_5: TI, area TI - PhG_L(R)_6_6: TH, area TH - pSTS_L(R)_2_1: rpSTS, rostroposterior superior temporal sulcus - pSTS_L(R)_2_2: cpSTS, caudoposterior superior temporal sulcus |
| Parietal (volume, thickness, surface area) | - SPL_L(R)_3_1: A7c, caudal area 7 - SPL_L(R)_3_2: A5l, lateral area 5 - SPL_L(R)_3_3: A7pc, postcentral area 7 - IPL_L(R)_6_1: A39c, caudal area 39 - IPL_L(R)_6_2: A39rd, rostrodorsal area 39 - IPL_L(R)_6_3: A40rd, rostrodorsal area 40 - IPL_L(R)_6_4: A40c, caudal area 40 - IPL_L(R)_6_5: A39rv, rostroventral area 39 - IPL_L(R)_6_6: A40rv, rostroventral area 40 - PCun_L(R)_4_1: A7m, medial area 7 - PCun_L(R)_4_2: A5m, medial area 5 - PCun_L(R)_4_3: dmPOS, dorsomedial parietooccipital sulcus - PCun_L(R)_4_4: A31, area 31 - PoG_L(R)_4_1: A1/2/3ulhf, area 1/2/3 - PoG_L(R)_4_2: A1/2/3tonIa, area 1/2/3 - PoG_L(R)_4_3: A2, area 2 - PoG_L(R)_4_4: A1/2/3tru, area 1/2/3 |
| Occipital (volume, thickness, surface area) | - MVOcC_L(R)_5_1: cLinG, caudal lingual gyrus - MVOcC_L(R)_5_2: rCunG, rostral cuneus gyrus - MVOcC_L(R)_5_3: cCunG, caudal cuneus gyrus - MVOcC_L(R)_5_4: rLinG, rostral lingual gyrus - MVOcC_L(R)_5_5: vmPOS, ventromedial parietooccipital sulcus - LOcC_L(R)_4_1: mOccG, middle occipital gyrus - LOcC_L(R)_4_2: V5/MT+, area V5/MT+ - LOcC_L(R)_4_3: OPC, occipital polar cortex - LOcC_L(R)_4_4: iOccG, inferior occipital gyrus - LOcC_L(R)_1_1: msOccG/lsOccG, medial superior occipital gyrus/lateral superior occipital gyrus |
| Insular (volume, thickness, surface area) | - Ins_L(R)_3_1: vIa, ventral agranular insula - Ins_L(R)_3_2: G/vId/vIg/dIg, hypergranular insula/ventral dysgranular and granular insula/dorsal granular insula - Ins_L(R)_3_3: dIa/dId, dorsal agranular insula/dorsal dysgranular insula |
| Limbic (volume, thickness, surface area) | - CG_L(R)_5_1: A23d, dorsal area 23 - CG_L(R)_5_2: A32p, pregenual area 32 - CG_L(R)_5_3: A23v, ventral area 23 - CG_L(R)_5_4: A23c, caudal area 23 - CG_L(R)_5_5: A32sg, subgenual area 32 |

**Supplementary Table 1.** Included parcels in each coarse area’s brain age model.

| **Models** | **Before correction** | | **After correction^1^** | |
| --- | --- | --- | --- | --- |
|  | MAE | r (p-value) | MAE | r (p-value) |
| **Male** | | | | |
| Whole brain | 5.98 | 0.34 (< 0.001) | 3.35 | 0.86 (< 0.001) |
| Subcortical | 6.25 | 0.23 (< 0.001) | 1.76 | 0.96 (< 0.001) |
| Frontal | 6.26 | 0.22 (< 0.001) | 2.96 | 0.91 (< 0.001) |
| Temporal | 6.48 | 0.08 (< 0.001) | 1.26 | 0.99 (< 0.001) |
| Parietal | 6.41 | 0.15 (< 0.001) | 0.93 | 0.99 (< 0.001) |
| Insula | 6.41 | 0.15 (< 0.001) | 0.72 | 0.99 (< 0.001) |
| Limbic | 6.41 | 0.15 (< 0.001) | 1.36 | 0.98 (< 0.001) |
| Occipital | 6.37 | 0.16 (< 0.001) | 1.13 | 0.98 (< 0.001) |
| **Female** | | | | |
| Whole brain | 6.19 | 0.31 (< 0.001) | 2.85 | 0.90 (< 0.001) |
| Subcortical | 6.38 | 0.25 (< 0.001) | 2.21 | 0.94 (< 0.001) |
| Frontal | 6.47 | 0.22 (< 0.001) | 2.46 | 0.93 (< 0.001) |
| Temporal | 6.73 | 0.05 (< 0.001) | 5.25 | 0.73 (< 0.001) |
| Parietal | 6.63 | 0.12 (< 0.001) | 1.75 | 0.98 (< 0.001) |
| Insula | 6.74 | 0.02 (< 0.001) | 2.87 | 0.95 (< 0.001) |
| Limbic | 6.65 | 0.09 (< 0.001) | 1.28 | 0.98 (< 0.001) |
| Occipital | 6.56 | 0.15 (< 0.001) | 1.52 | 0.97 (< 0.001) |

**Supplementary Table 2.** Male and female models’ performance on their respective test sets, before and after age-bias correction. ^1^: Statistics after correction as described by Beheshti et al. (2019). MAE: Mean absolute error. r: Pearson’s correlation coefficient.

| **Regions** | **AD** | | | **ADHD** | | | | **CD** | | | | **DD** | | | | **OCD** | | | | **ODD** | | | | **PSYPATHO** | | | |
| --- | --- | --- | --- | --- | --- | --- | --- | --- | --- | --- | --- | --- | --- | --- | --- | --- | --- | --- | --- | --- | --- | --- | --- | --- | --- | --- | --- |
|  | ***d*** | ***p_fdr_*** | | ***d*** | | ***p_fdr_*** | | ***d*** | | ***p_fdr_*** | | ***d*** | | ***p_fdr_*** | | ***d*** | | ***p_fdr_*** | | ***d*** | | ***p_fdr_*** | | ***d*** | | ***p_fdr_*** | |
| **Male** | | | | | | | | | | | | | | | | | | | | | | | | | | | |
| Whole Brain | -0.05 | 0.761 | | -0.07 | | 0.263 | | -0.01 | | 0.907 | | -0.15 | | 0.785 | | -0.07 | | 0.533 | | -0.02 | | 0.940 | | -0.05 | | 0.317 | |
| Frontal | -0.05 | 0.761 | | -0.09 | | 0.130 | | -0.09 | | 0.518 | | -0.21 | | 0.785 | | -0.04 | | 0.800 | | -0.04 | | 0.940 | | -0.06 | | 0.301 | |
| Temporal | 0.11 | 0.160 | | 0.00 | | 0.942 | | 0.15 | | 0.518 | | 0.04 | | 0.963 | | -0.01 | | 0.928 | | -0.01 | | 0.940 | | 0.01 | | 0.860 | |
| Parietal | 0.11 | 0.160 | | 0.00 | | 0.942 | | 0.09 | | 0.518 | | 0.01 | | 0.963 | | 0.05 | | 0.777 | | -0.06 | | 0.889 | | 0.01 | | 0.818 | |
| Occipital | -0.01 | 0.905 | | -0.06 | | 0.301 | | 0.06 | | 0.578 | | -0.30 | | 0.785 | | 0.00 | | 0.928 | | -0.09 | | 0.833 | | -0.03 | | 0.818 | |
| Subcortical | -0.03 | 0.761 | | -0.09 | | 0.130 | | -0.11 | | 0.518 | | -0.20 | | 0.785 | | -0.14 | | 0.095 | | -0.08 | | 0.833 | | -0.08 | | 0.110 | |
| Limbic | 0.00 | 0.990 | | -0.03 | | 0.632 | | 0.02 | | 0.907 | | 0.12 | | 0.785 | | -0.02 | | 0.928 | | -0.02 | | 0.940 | | -0.02 | | 0.818 | |
| Insula | -0.03 | 0.761 | | 0.02 | | 0.716 | | 0.08 | | 0.536 | | -0.13 | | 0.785 | | 0.11 | | 0.208 | | 0.00 | | 0.940 | | 0.01 | | 0.818 | |
| **Female** | | | | | | | | | | | | | | | | | | | | | | | | | | | |
| Whole Brain | 0.05 | | 0.876 | | -0.05 | | 0.602 | | -0.18 | | 0.336 | | 0.04 | | 0.886 | | -0.02 | | 0.822 | | -0.12 | | 0.808 | | 0.01 | | 0.867 |
| Frontal | 0.01 | | 0.894 | | -0.07 | | 0.602 | | -0.09 | | 0.531 | | -0.15 | | 0.859 | | -0.12 | | 0.448 | | -0.09 | | 0.808 | | -0.03 | | 0.867 |
| Temporal | 0.06 | | 0.876 | | 0.05 | | 0.602 | | 0.20 | | 0.336 | | 0.57 | | 0.302 | | 0.10 | | 0.448 | | 0.04 | | 0.833 | | 0.06 | | 0.740 |
| Parietal | 0.01 | | 0.894 | | 0.06 | | 0.602 | | -0.19 | | 0.336 | | -0.09 | | 0.859 | | 0.02 | | 0.822 | | -0.08 | | 0.808 | | 0.04 | | 0.867 |
| Occipital | 0.07 | | 0.876 | | 0.00 | | 0.938 | | -0.09 | | 0.531 | | -0.26 | | 0.859 | | -0.02 | | 0.822 | | -0.05 | | 0.833 | | 0.02 | | 0.867 |
| Subcortical | 0.03 | | 0.876 | | -0.14 | | 0.067 | | -0.11 | | 0.531 | | 0.10 | | 0.859 | | 0.04 | | 0.822 | | -0.02 | | 0.837 | | -0.01 | | 0.867 |
| Limbic | 0.04 | | 0.876 | | -0.03 | | 0.740 | | 0.03 | | 0.780 | | 0.16 | | 0.859 | | -0.07 | | 0.695 | | -0.05 | | 0.833 | | -0.01 | | 0.867 |
| Insula | -0.01 | | 0.894 | | 0.00 | | 0.938 | | 0.12 | | 0.531 | | 0.32 | | 0.859 | | -0.02 | | 0.822 | | -0.02 | | 0.837 | | 0.00 | | 0.992 |

**Supplementary Table 3.** Clinical samples group effects for each brain region stratified by BAG derived from the male and female-only models. *d*: Cohen’s *d* effect size. *p_fdr_*: FDR-corrected p-value (< 0.05 are **bolded**). AD: Anxiety disorder. ADHD: Attention-deficit/hyperactivity disorder. CD: Conduct disorder. OCD: Obsessive-compulsive disorder. ODD: Oppositional defiant disorder. DD: Depressive disorder. PSYPATHO: Participants with at least one psychiatric disorder.

| **Regions** | **MC/HSI** | | **MC/HE** | | **HC/LB** | | **LC/LB** | |
| --- | --- | --- | --- | --- | --- | --- | --- | --- |
|  | ***r_partial_*** | ***p_fdr_*** | ***r_partial_*** | ***p_fdr_*** | ***r_partial_*** | ***p_fdr_*** | ***r_partial_*** | ***p_fdr_*** |
| **Male** | | | | | | | | |
| Whole Brain | -0.02 | 0.611 | -0.02 | 0.411 | 0.03 | 0.069 | -0.01 | 0.811 |
| Frontal | 0.00 | 0.852 | -0.03 | 0.411 | 0.04 | 0.069 | 0.00 | 0.934 |
| Temporal | 0.00 | 0.852 | 0.01 | 0.425 | -0.02 | 0.422 | 0.01 | 0.811 |
| Parietal | 0.00 | 0.852 | -0.02 | 0.411 | -0.03 | 0.069 | 0.02 | 0.811 |
| Occipital | -0.03 | 0.345 | -0.02 | 0.411 | 0.01 | 0.644 | 0.01 | 0.811 |
| Subcortical | -0.02 | 0.627 | -0.02 | 0.411 | 0.03 | 0.075 | 0.00 | 0.934 |
| Limbic | -0.03 | 0.345 | -0.02 | 0.411 | 0.00 | 0.991 | 0.01 | 0.811 |
| Insula | 0.00 | 0.852 | -0.01 | 0.703 | -0.03 | 0.075 | -0.01 | 0.933 |
| **Female** | | | | | | | | |
| Whole Brain | 0.00 | 0.895 | -0.02 | 0.461 | 0.03 | 0.154 | -0.02 | 0.465 |
| Frontal | 0.00 | 0.895 | -0.02 | 0.461 | 0.01 | 0.637 | 0.00 | 0.941 |
| Temporal | 0.00 | 0.895 | -0.02 | 0.461 | 0.01 | 0.732 | 0.01 | 0.941 |
| Parietal | -0.02 | 0.839 | 0.00 | 0.975 | -0.03 | 0.211 | 0.03 | 0.377 |
| Occipital | -0.02 | 0.839 | 0.01 | 0.798 | 0.00 | 0.937 | 0.02 | 0.465 |
| Subcortical | 0.00 | 0.895 | -0.02 | 0.531 | 0.03 | 0.154 | 0.00 | 0.941 |
| Limbic | -0.01 | 0.839 | -0.01 | 0.798 | 0.03 | 0.154 | 0.00 | 0.941 |
| Insula | 0.02 | 0.839 | 0.03 | 0.461 | 0.01 | 0.732 | -0.01 | 0.941 |

**Supplementary Table 4.** Partial correlation coefficient (*r_partial_*) between BAG values per regions derived from the male and female only models and the membership values for each fuzzy cognitive and behavioral profile. *p_fdr_*: FDR-corrected p-value (< 0.05 are **bolded**). MC/HSI: Moderate cognition and high stress/internalization. MC/HE: Moderate cognition and high externalization. HC/LB: High cognition and low behavior. LC/LB: Low cognition and low behavior.

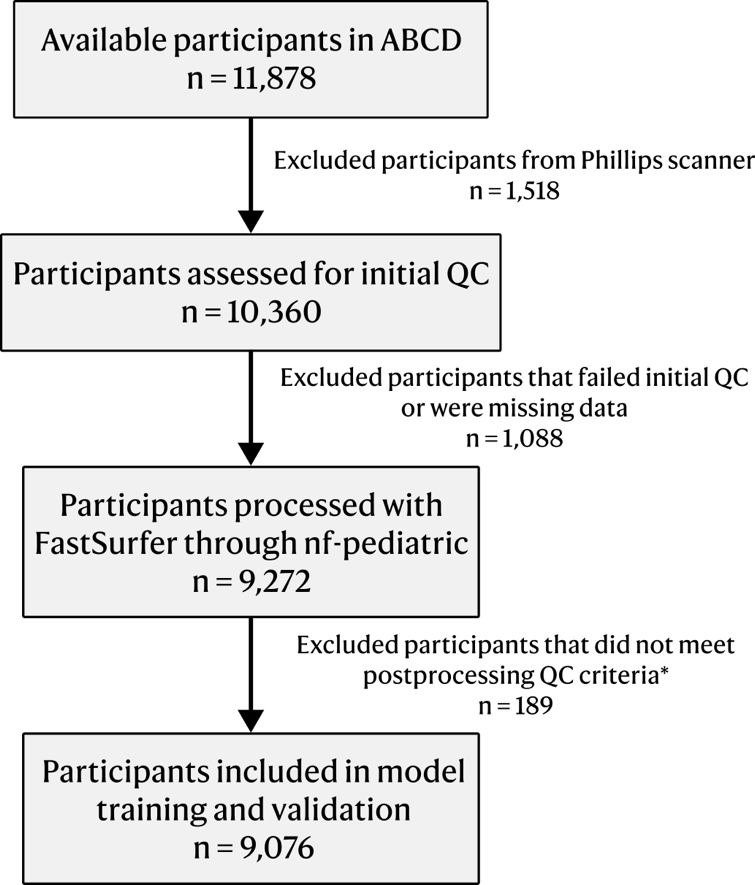
**Supplementary Figure 1.** Flowchart of participants’ exclusion. *: Participants who were either 1) missing cortical/subcortical segmentations or 2) had metric values higher or lower than five times the interquartile range ($\pm5 \times IQR$) in a single parcel.

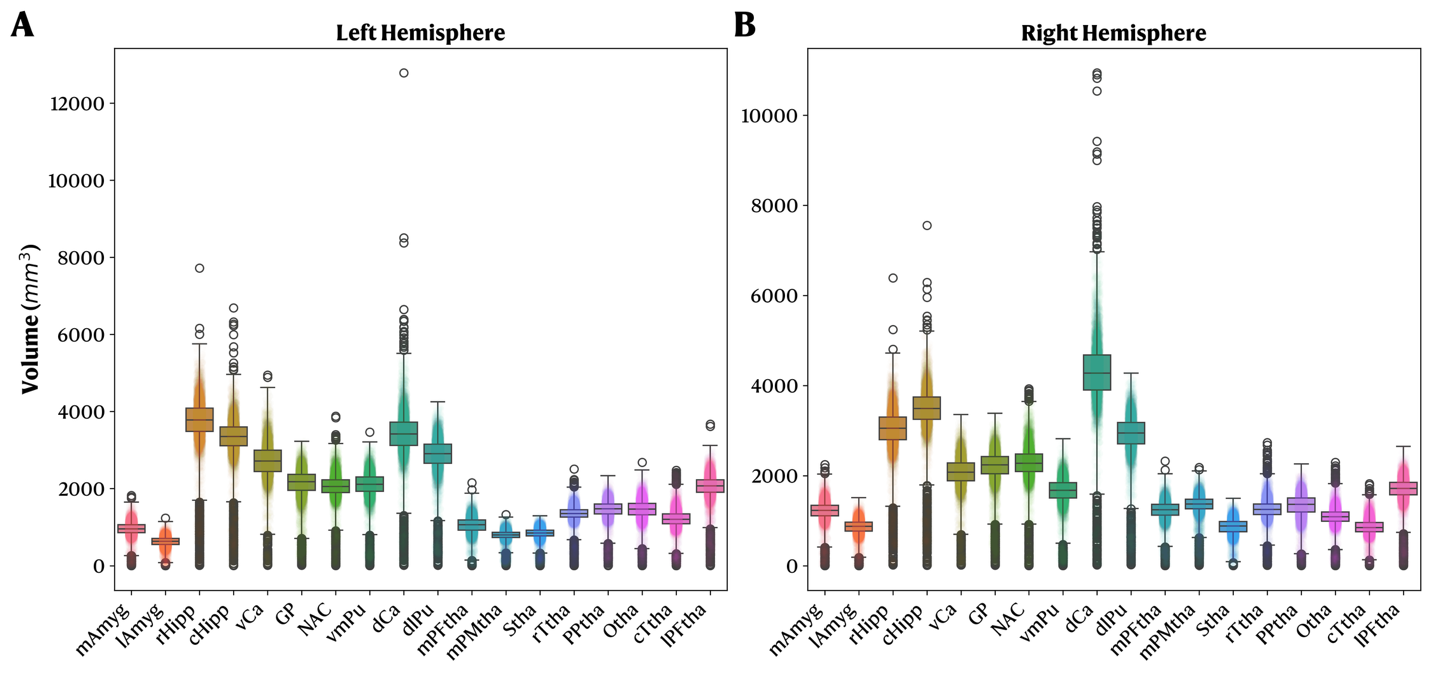
**Supplementary Figure 2.** Distribution of subcortical volumes for left and right hemispheres before outlier removal. Whiskers denote the $3\times IQR$. **A.** Left Hemisphere. **B.** Right Hemisphere.

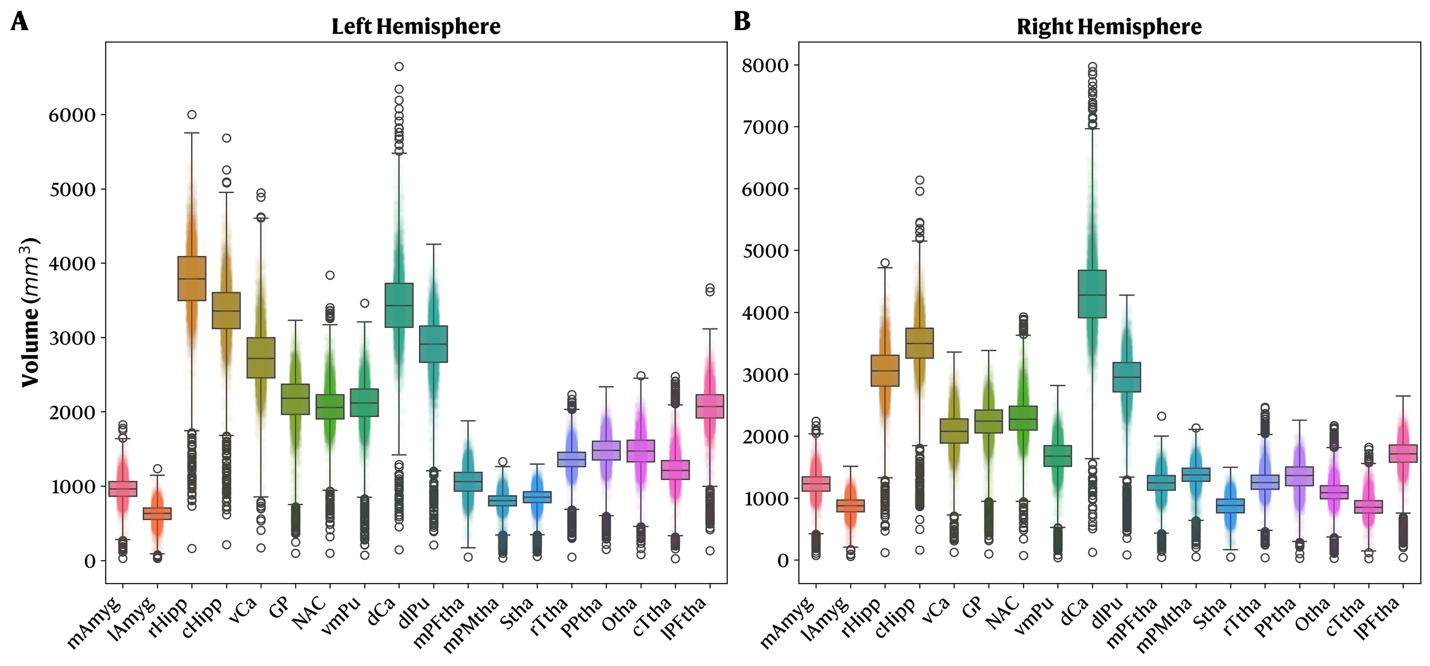
**Supplementary Figure 3.** Distribution of subcortical volumes for left and right hemispheres after outlier removal. Whiskers denote the $3\times IQR$. **A.** Left Hemisphere. **B.** Right Hemisphere.

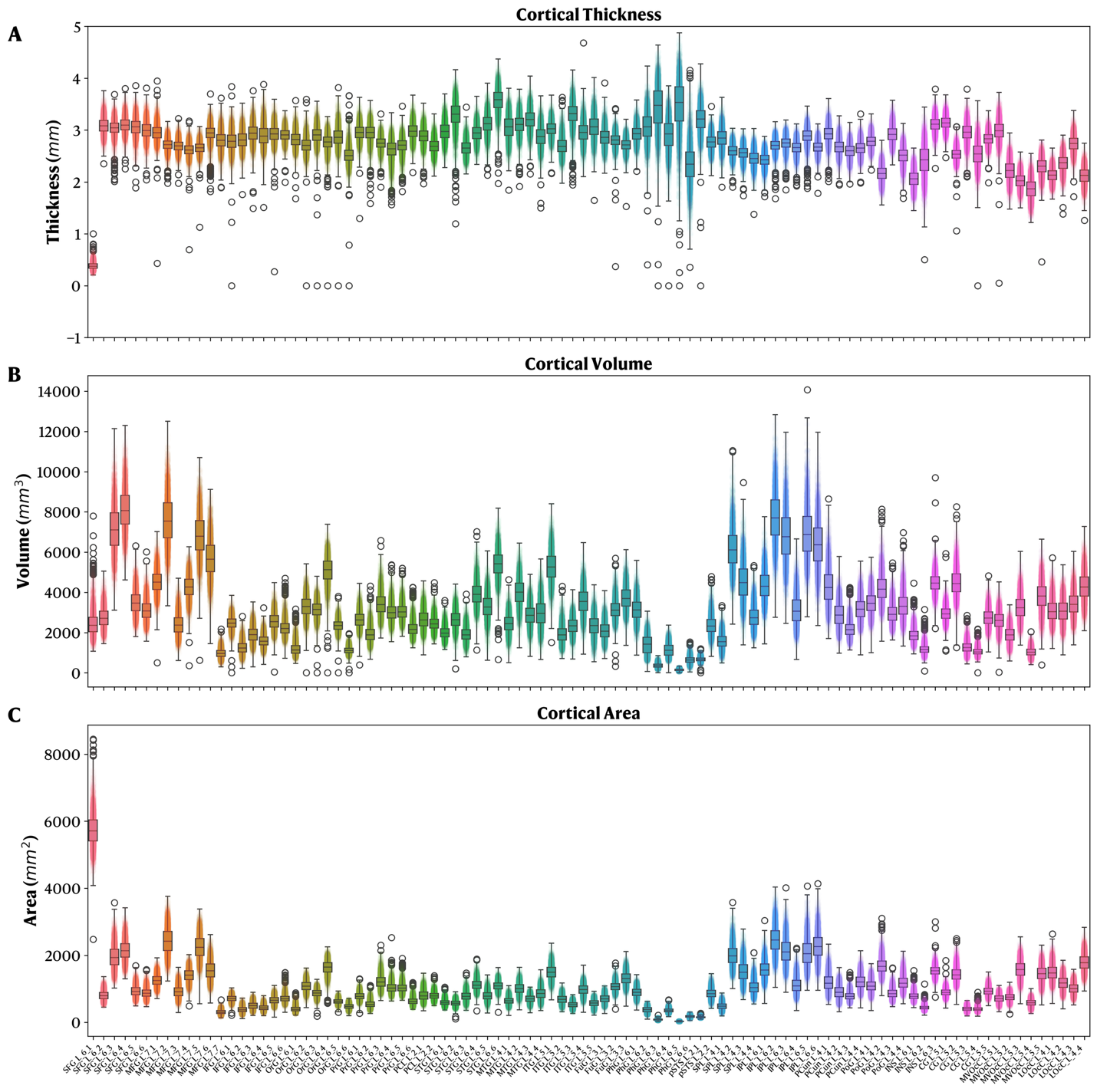
**Supplementary Figure 4.** Distribution of cortical thickness, volume, and surface area for the **left hemisphere** before outlier removal. Whiskers denote the $3\times IQR$. **A.** Thickness. **B.** Volume. **C.** Surface Area.

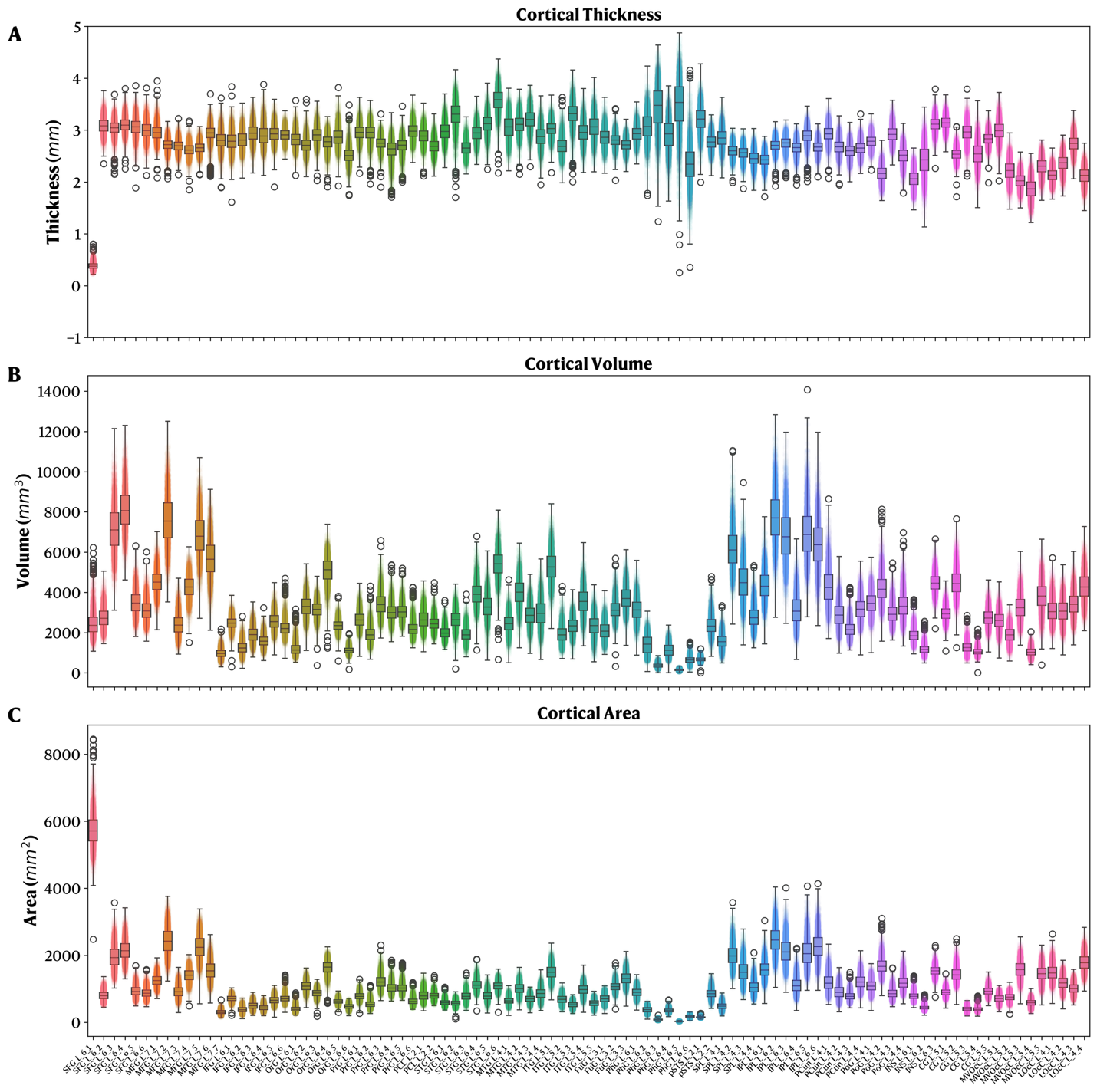
**Supplementary Figure 5.** Distribution of cortical thickness, volume, and surface area for the **left hemisphere** after outlier removal. Whiskers denote the $3\times IQR$. **A.** Thickness. **B.** Volume. **C.** Surface Area.

**Supplementary Figure 6.** Distribution of cortical thickness, volume, and surface area for the **right hemisphere** before outlier removal. Whiskers denote the $3\times IQR$. **A.** Thickness. **B.** Volume. **C.** Surface Area
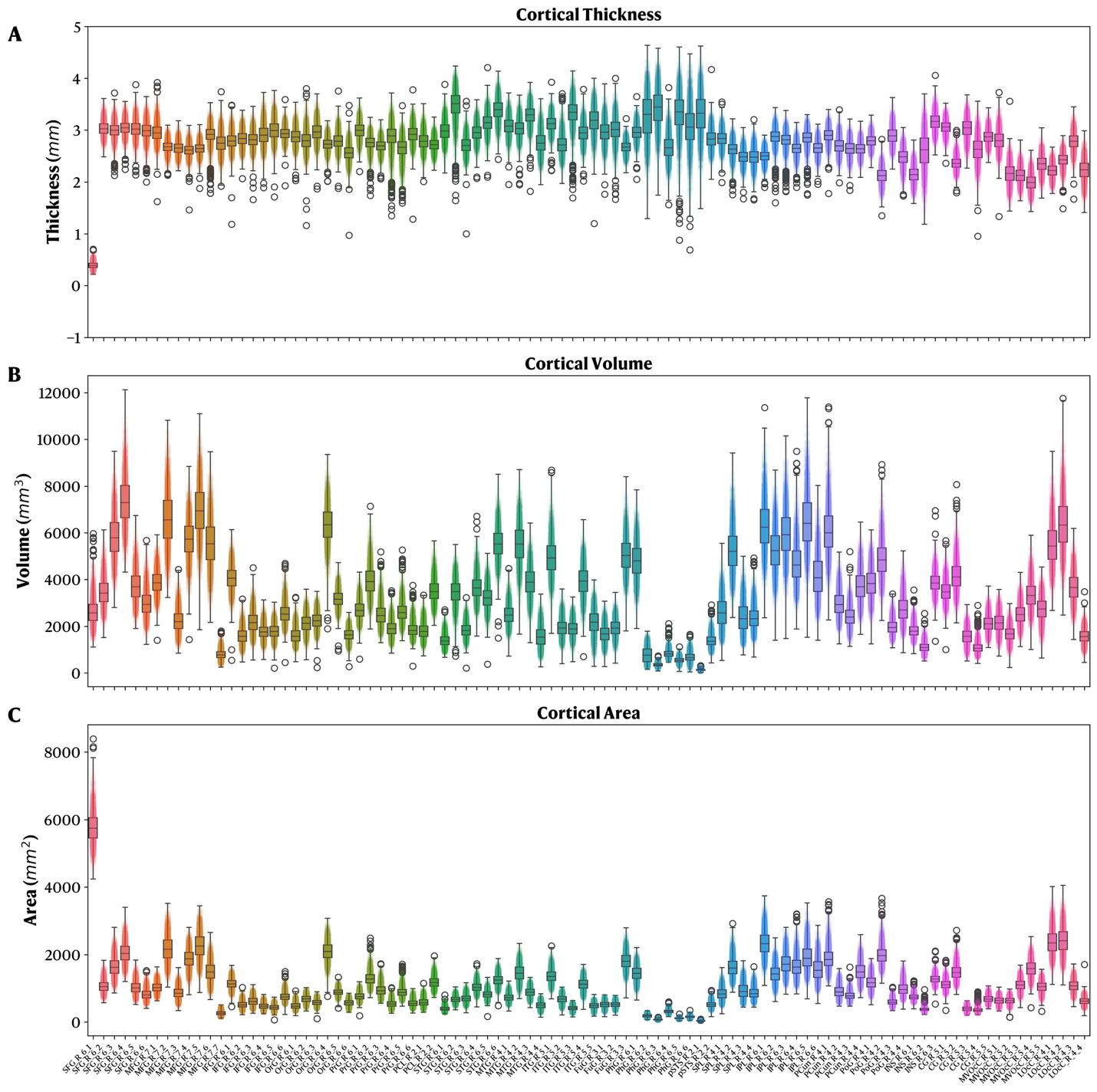
**.**

**Supplementary Figure 7.** Distribution of cortical thickness, volume, and surface area for the **right hemisphere** after outlier removal. Whiskers denote the $3\times IQR$. **A.** Thickness. **B.** Volume. **C.** Surface Area
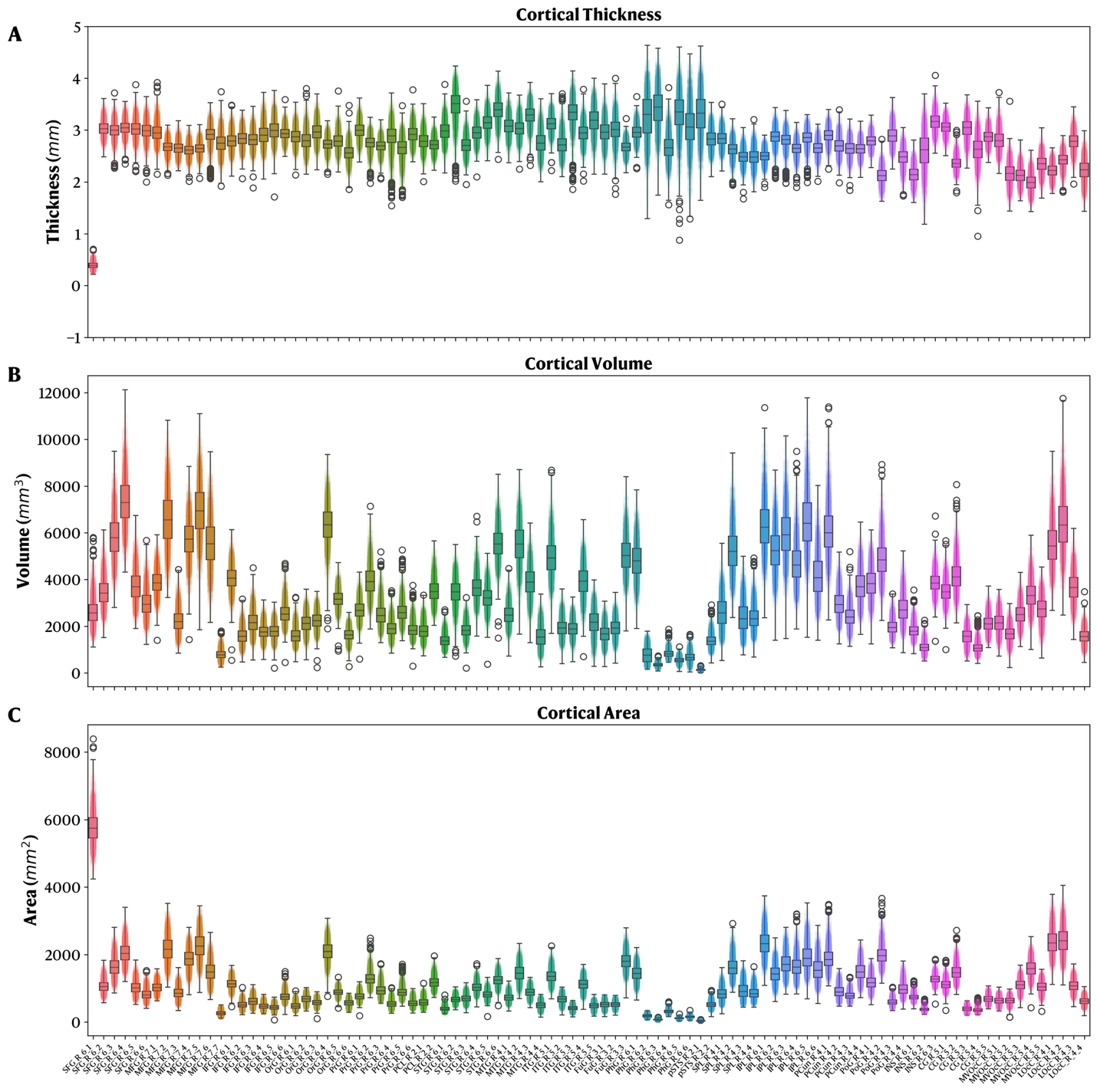
.

**Supplementary** **Figure 8.** Overview of the participants’ site distribution, in addition to the characteristics of the reference site used as a target for data harmonization (site 16). **A.** Participant distribution across sites in percentage. **B.** Sex distribution within the reference site (site 16). **C.** Age distribution within the reference site (site 16).
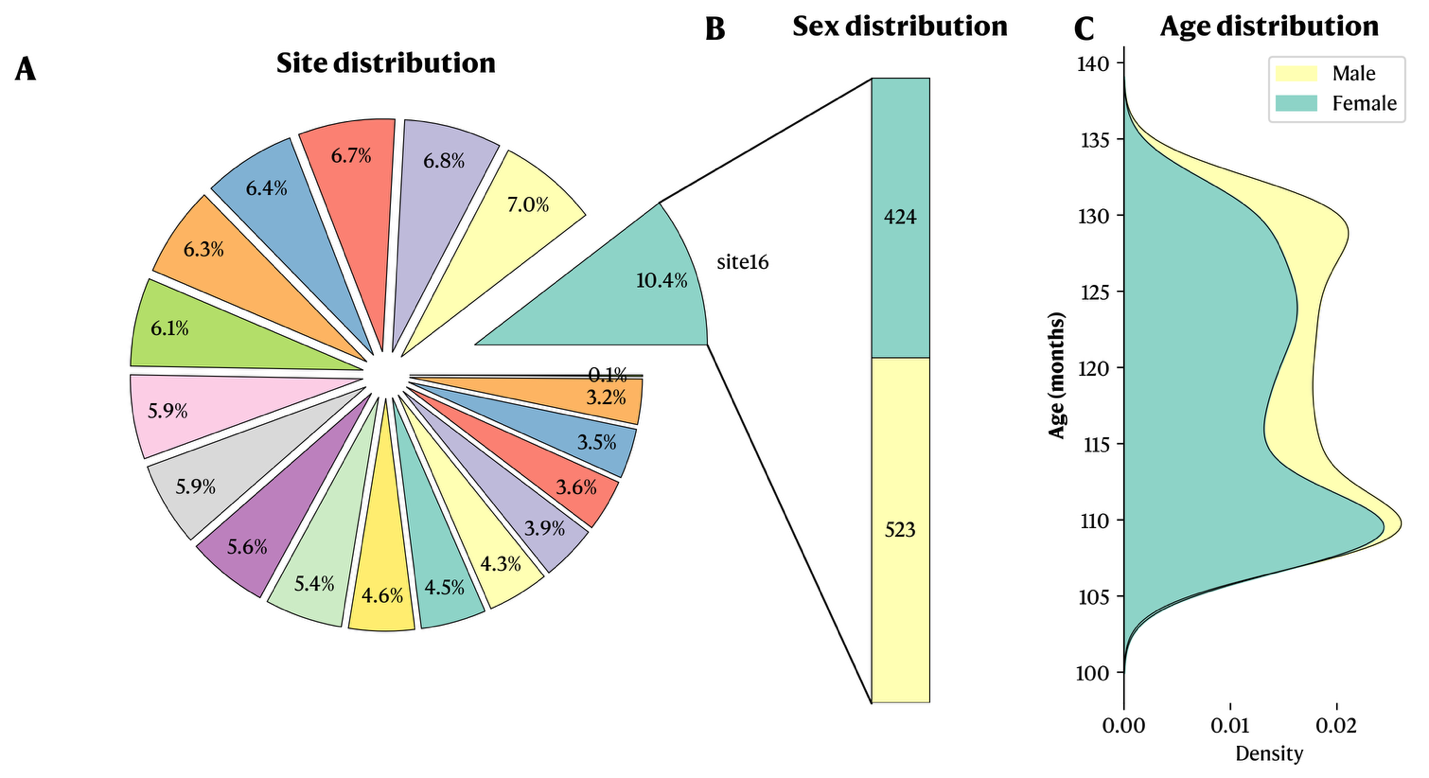

**Supplementary Figure 9.**
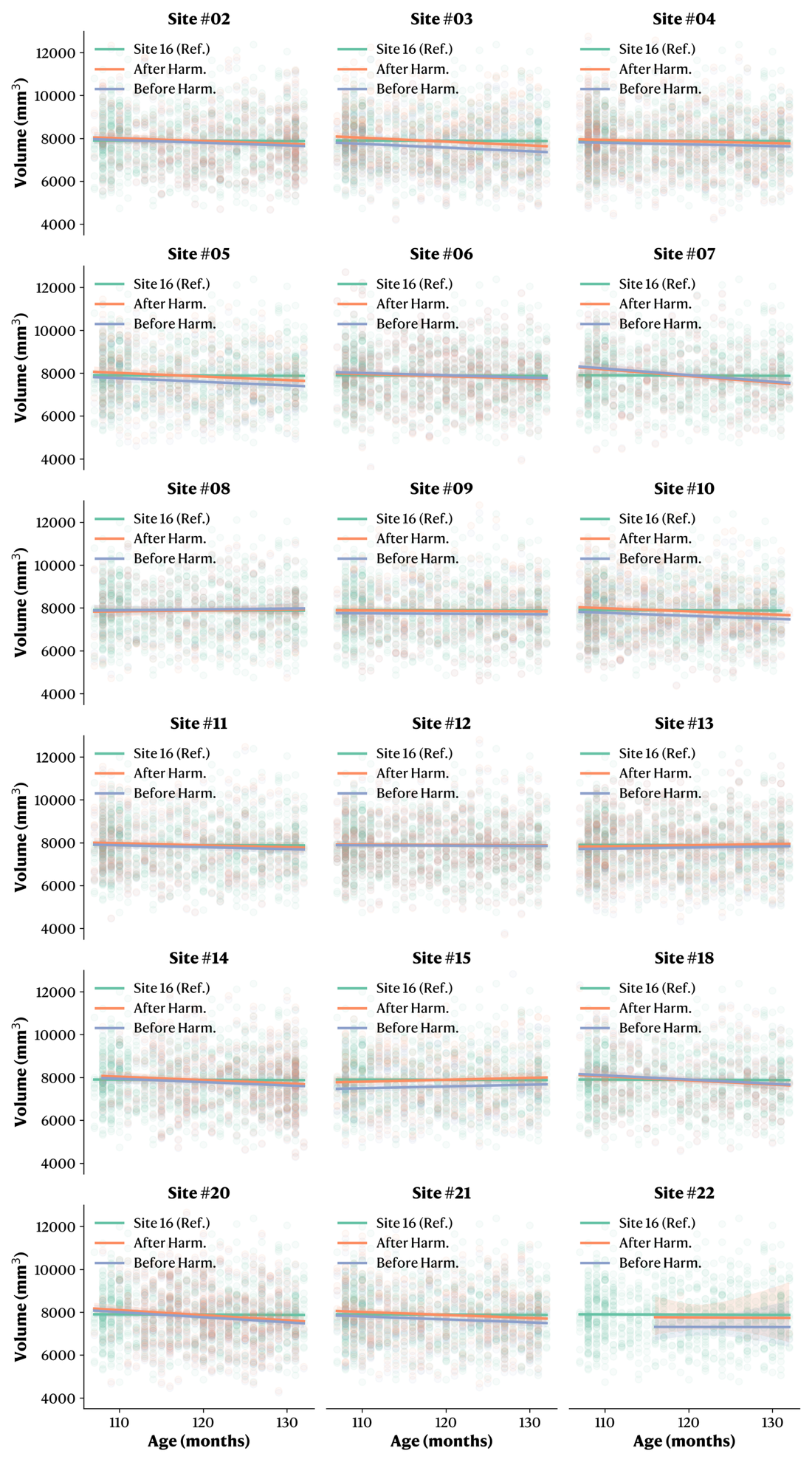
 Example visualization of the pairwise harmonization to the reference site (site 16) for the volume metric in the inferior parietal lobule A39rd region. The green curve represents the reference site (site 16), while the blue curve is the moving site before harmonization, and the orange curve represents the moving site after harmonization.

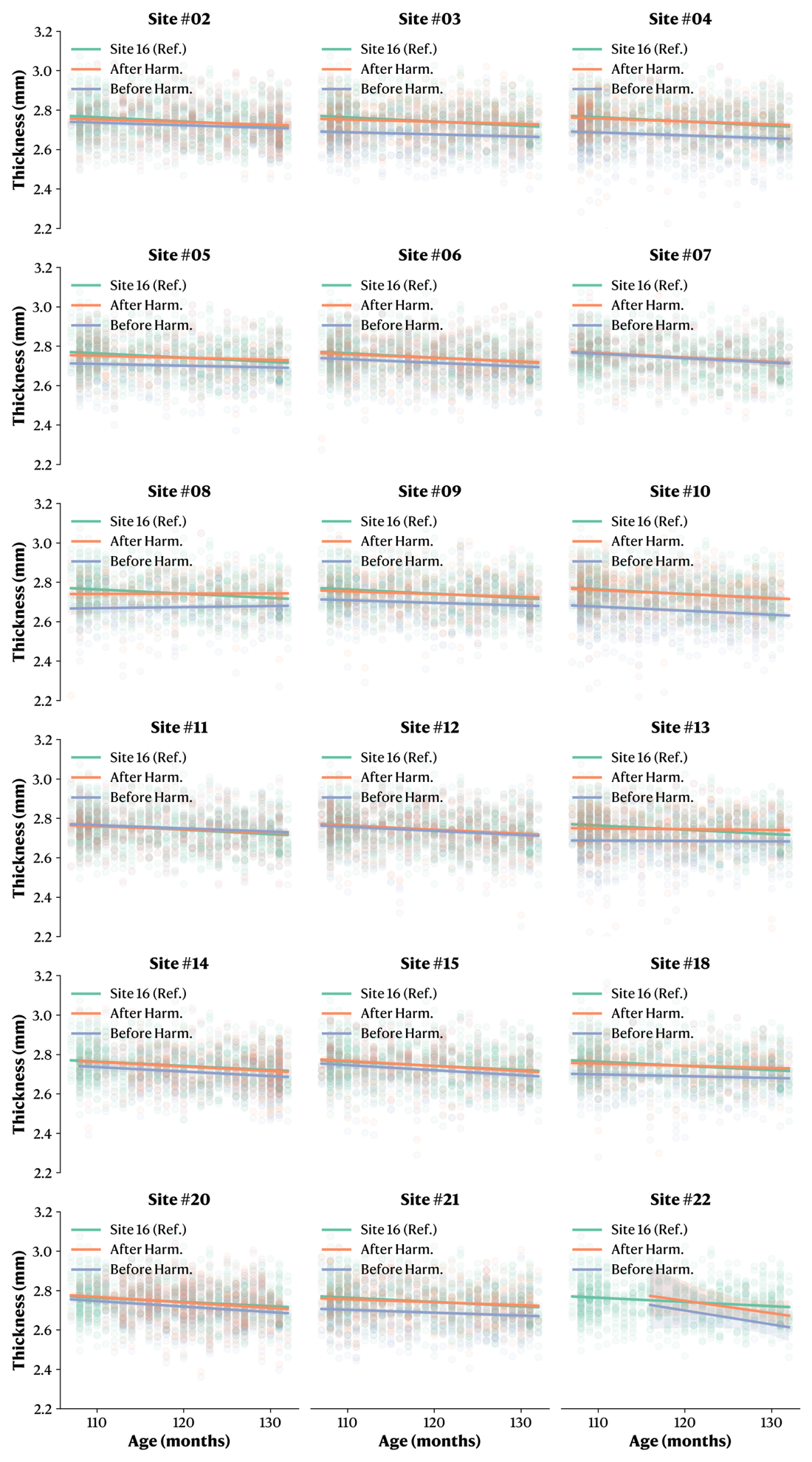
**Supplementary Figure 10.**  Example visualization of the pairwise harmonization to the reference site (site 16) for the cortical thickness metric in the inferior parietal lobule A39rd region. The green curve represents the reference site (site 16), while the blue curve is the moving site before harmonization, and the orange curve represents the moving site after harmonization.

**Supplementary Figure 11.**  Example visualization of the pairwise harmonization to the reference site (site 16) for the surface area metric in the inferior parietal lobule A39rd region. The green curve represents the reference site (site 16), while the blue curve is the moving site before harmonization, and the orange curve represents the moving site after harmonization
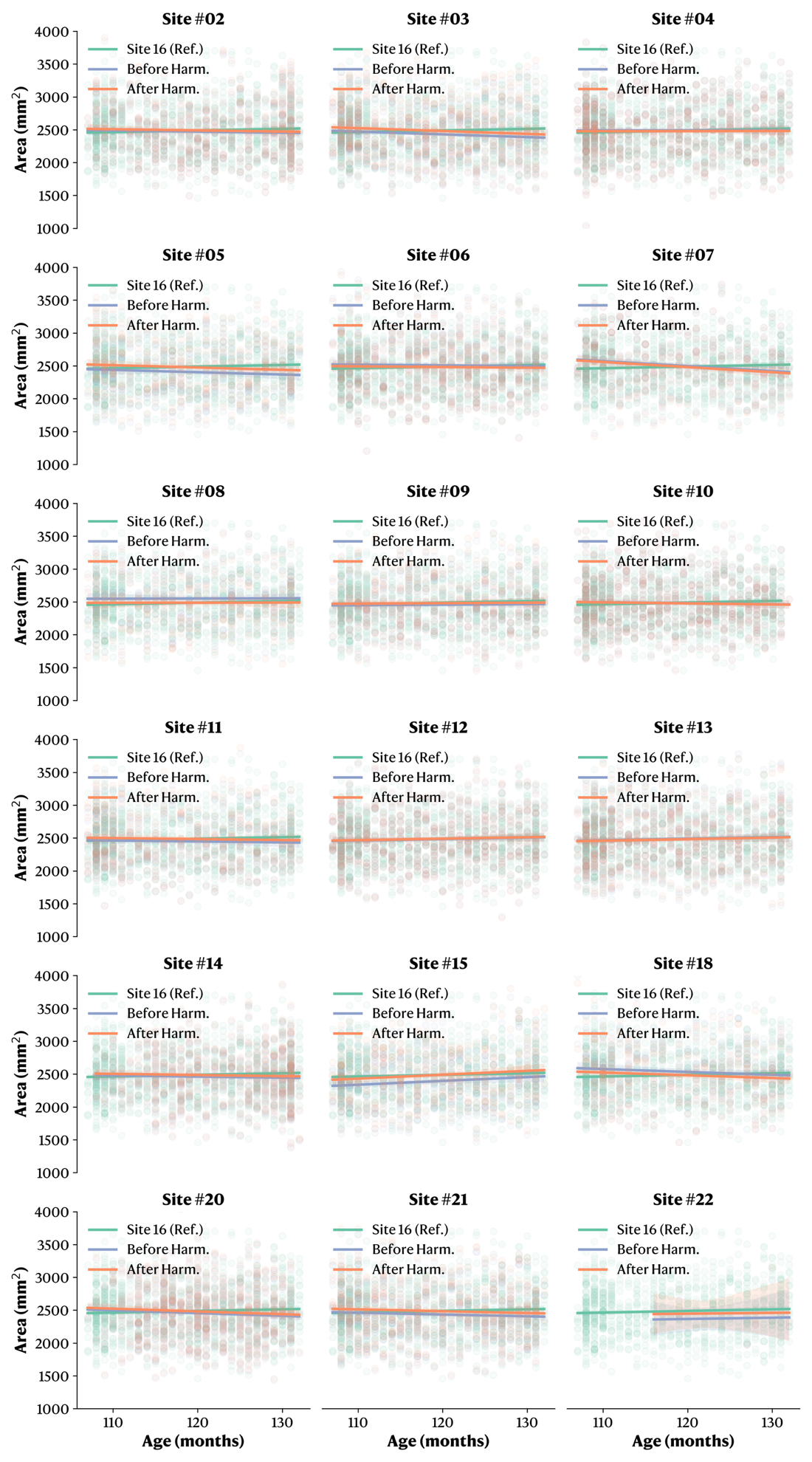
.

**Supplementary Figure 12.**
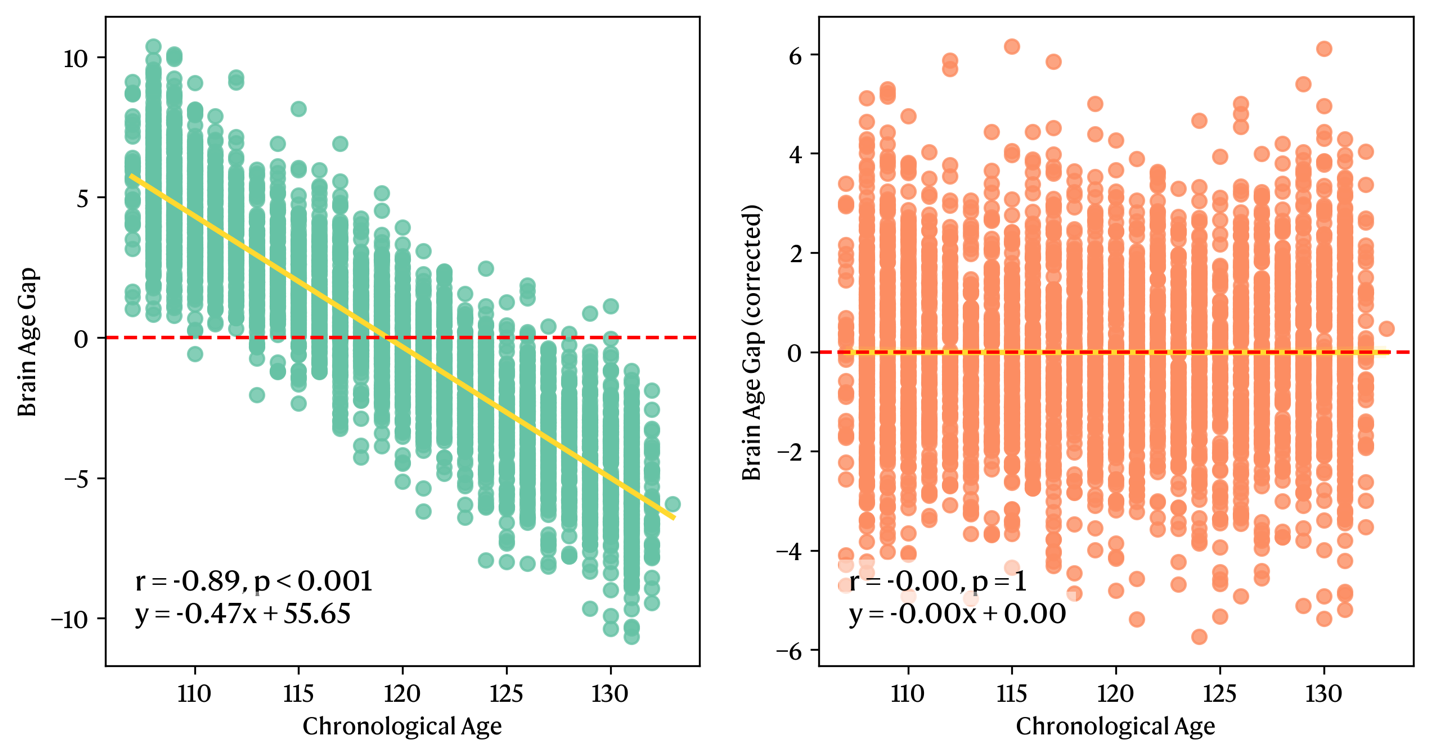
Visualization of the age bias in brain age prediction for the whole-brain model before and after correction. The applied correction was developed and described by Beheshti *et al.* (2019)^1^.

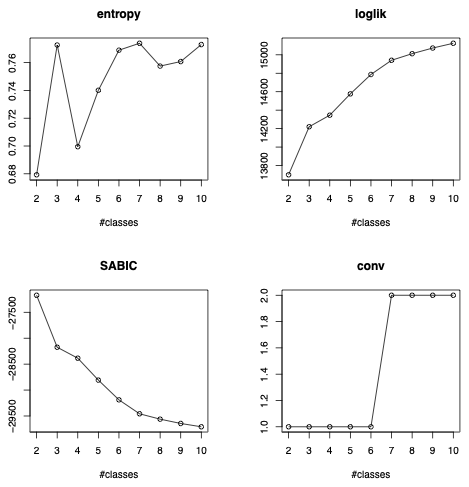
**Supplementary Figure 13.** Statistical criteria plots assessing the fit of multiple latent class models (2 to 10). The final selection was based on the entropy and convergence status. Loglik: Log-likelihood. SABIC: Sample-adjusted Bayesian Information Criteria. Conv: Convergence status (1: converged and 2: did not converge).
